# The effects of multi-day rTMS and cardiorespiratory fitness on working memory and local GABA concentration

**DOI:** 10.1101/2020.09.09.288910

**Authors:** Joshua Hendrikse, Sarah Thompson, Chao Suo, Murat Yücel, Nigel C. Rogasch, James P. Coxon

**Affiliations:** Turner Institute for Brain and Mental Health and School of Psychological Sciences, Monash University, Melbourne, Victoria, Australia; Hopwood Centre for Neurobiology, Lifelong Health Theme, South Australian Health and Medical Research Institute (SAHMRI); Discipline of Psychiatry, Adelaide Medical School, University of Adelaide

**Keywords:** rTMS, metabolite, spectroscopy, GABA, working memory, neuroplasticity

## Abstract

Working memory (WM) refers to the capacity to temporarily retain and manipulate finite amounts of information; a critical process in complex behaviours such as reasoning, comprehension, and learning. This cognitive function is supported by a parietal-prefrontal network and linked to the activity of key neurotransmitters, such as gamma-aminobutyric acid (GABA). Impairments in WM are seen in a range of psychiatric and neurological disorders, and there are currently no effective treatments. In this study, we analysed secondary outcome measures from a trial investigating the effects of multi-day rTMS on cognition. Participants received four days of 20 Hz rTMS to an individualised region of left parietal cortex in one week, and an individualised region of pre-supplementary motor area (pre-SMA) in a separate week. We assessed changes to WM function before and after each week of stimulation (N = 39), and changes to GABA concentration before and after stimulation in week one using MR spectroscopy (N = 18 per stimulation condition). We hypothesised that parietal rTMS would enhance WM and alter GABA concentration at the site of stimulation, but this was not observed. Instead, we report some evidence of improved WM function following the first week of pre-SMA rTMS stimulation, and a generalised increase in GABA concentration across both parietal and pre-SMA voxels following pre-SMA rTMS. Additionally, we found that higher cardiorespiratory fitness was associated with greater WM improvement following pre-SMA stimulation. This study does not support the use of parietal multi-day rTMS for the enhancement of working memory. In contrast, the results suggest that increasing cardiorespiratory fitness may provide a novel approach to enhance the effects of pre-SMA rTMS on cognition.

## Introduction

Working memory (WM) refers to the ability to temporarily retain and manipulate finite amounts of information (Baddeley, 2010). The importance of this cognitive process is underscored by its role in complex behaviours such as reasoning, comprehension, and learning, and its association with overall intelligence (Baddeley, 1992). Furthermore, WM impairments are common across a wide range of psychiatric and neurological disorders, including schizophrenia (Silver, Feldman, Bilker, & Gur, 2003) and Alzheimer’s disease (Baddeley, Bressi, Della Sala, Logie, & Spinnler, 1991). Despite the near ubiquitous nature of WM deficits across brain disorders, there are currently no effective methods for improving WM. Current therapies (e.g. pharmacological and cognitive remediation strategies) are associated with small/moderate effect sizes (Lett, Voineskos, Kennedy, Levine, & Daskalakis, 2014), and are therefore of limited clinical value. Thus, there is a need to investigate novel approaches to alleviate WM deficits.

Repetitive transcranial magnetic stimulation (rTMS) is a non-invasive brain stimulation method with the capacity to induce neuroplasticity in targeted cortical regions (Hallett, 2007) and alter working memory performance. For example, a single dose of rTMS to the dorsolateral prefrontal cortex (dlPFC) can induce transient improvements in WM (Brunoni & Vanderhasselt, 2014; Hoy et al., 2016). However, past trials examining the effects of multi-day dlPFC rTMS (i.e. repeating rTMS sessions over multiple days) on WM have shown mixed results. Some studies have showed WM improvement in healthy (Bagherzadeh, Khorrami, Zarrindast, Shariat, & Pantazis, 2016) and patient populations (Barr et al., 2011), whereas others failed to demonstrate changes in WM in either healthy or patient cohorts (Gaudeau-Bosma et al., 2013; Guse et al., 2013). Overall, meta-analysis has shown a significant effect of high-frequency dlPFC rTMS on WM, although the effect size is small potentially limiting clinical utility (Brunoni & Vanderhasselt, 2014).

Targeting other cortical regions within the WM network with rTMS may also prove beneficial. Several studies have shown transient improvements in WM following a single stimulation session targeting parietal cortex (Albouy, Weiss, Baillet, & Zatorre, 2017; Hamidi, Tononi, & Postle, 2008; Luber et al., 2007b; Yamanaka, Yamagata, Tomioka, Kawasaki, & Mimura, 2010). Sustained improvement in associative memory function has also been reported following 5 sessions rTMS to a region of the inferior parietal cortex with strong functional connectivity to the hippocampus (Wang et al., 2014; Wang & Voss, 2015), although these findings have not always replicated (Hendrikse et al., 2020). In addition to dlPFC and parietal cortex, the n-back WM task also engages secondary motor regions (e.g. pre-supplementary motor area (pre-SMA) and striatum) (Buchsbaum & D’Esposito, 2019; Rottschy et al., 2012), which may reflect motor preparation/initiation (Buchsbaum & D’Esposito, 2019), the encoding of covert motor ‘traces’ for maintenance of WM content (Marvel, Morgan, & Kronemer, 2019), or the prevention of interference during WM (Irlbacher, Kraft, Kehrer, & Brandt, 2014). Overall, the parietal cortex and pre-SMA may provide effective alternative targets to improve WM. However, the effects of multi-day rTMS on these regions is not yet known.

We know that a single session of rTMS is capable of modulating GABA-mediated neurotransmission (Fitzgerald, Fountain, & Daskalakis, 2006), but the neurophysiological effects of multi-day rTMS are not well understood. Studies have used magnetic resonance spectroscopy (MRS), which provides an estimate of the total concentration of GABA (Stagg, Bachtiar, & Johansen-berg, 2011), to show that a single dose of high-frequency rTMS to primary motor cortex (Stagg et al., 2009) and inferior parietal cortex (Vidal-Piñeiro et al., 2015) can increase GABA. However, it remains unclear whether multi-day rTMS can induce long-lasting changes in GABA concentration at the site of stimulation. Given that GABA concentration is associated with WM performance (Michels et al., 2012; Yoon, Grandelis, & Maddock, 2016), increasing GABA concentrations may be an important candidate mechanism for improving WM with rTMS.

Cardiorespiratory exercise is another factor known to influence neuroplasticity, and may provide an effective method for augmenting the effects of rTMS (Cirillo, Lavender, Ridding, & Semmler, 2009; Hendrikse, Kandola, Coxon, Rogasch, & Yücel, 2017; Andrews et al., 2019). In particular, high-intensity exercise is known to influence the concentration of GABA in the brain (Coxon et al., 2018; Maddock, Casazza, Fernandez, & Maddock, 2016;), and upregulate a host of mechanisms which influence neuroplasticity (e.g. increased cerebral blood flow, BDNF and neurotrophin circulation; see Kandola, Hendrikse, Lucassen, & Yücel, 2016; Voss, Vivar, Kramer, & van Praag, 2013). Furthermore, cardiorespiratory fitness level (Ishihara, Miyazaki, Tanaka, & Matsuda, 2020), and both acute (Pontifex, Hillman, Fernhall, Thompson, & Valentini, 2009) and sustained regular exercise (Guiney & Machado, 2013) have also been associated with enhanced WM function. However, it remains unknown whether cardiorespiratory fitness, and/or regular exercise, influence the effects of multi-day rTMS.

In summary, multi-day rTMS studies targeting prefrontal cortex have had mixed results on WM, and it is unclear whether targeting other regions may improve WM. It is also unclear whether multi-day rTMS can result in long-lasting changes in neurotransmitter concentrations, such as GABA, or if behavioural and neurophysiological effects are modulated by cardiorespiratory fitness. In this study, we analysed secondary outcome measures from a trial investigating the effects of multi-day 20 Hz rTMS to parietal cortex and pre-SMA on cognitive function (Hendrikse et al., 2020). The aims of this study were threefold. The first aim was to assess whether multi-day 20 Hz rTMS can induce long-lasting improvements in WM. The second aim was to assess whether multi-day rTMS can alter local GABA concentrations at the site of stimulation. The third aim was to investigate whether cardiorespiratory fitness influenced the magnitude of rTMS-induced effects. We expected multi-day rTMS would improve WM performance, and that this would manifest following stimulation of both parietal cortex and pre-SMA. Further, we expected a significant change in GABA concentration specific to the stimulation site following multi-day rTMS. We also expected that the effects of multi-day rTMS on both WM and GABA concentration would be greater in individuals engaging in high levels of exercise, with higher cardiorespiratory fitness levels (CRF; i.e. VO_2_max score).

## Methods

### Ethics approval

This study was approved by the Monash University Human Research Ethics Committee and all subjects provided informed consent. The study conformed to the standards set by the Declaration of Helsinki. Participants were remunerated $100 for their participation.

### Participants

Participants were 40 right-handed healthy adults (52.5% female), aged 25.48 ± 9.35 years (mean ± SD; range 18 – 55), who were medication-free, and reported no contraindications to magnetic resonance imaging or transcranial magnetic stimulation and no history of psychological or neurological disorders.

To investigate the influence of exercise on the response to multi-day rTMS, we recruited and selected participants on the basis of self-reported weekly exercise levels. Participants were categorised into either a high physical activity group (high PA) or low physical activity group (low PA). The high PA group included individuals engaging in a minimum of 150 minutes of deliberate exercise each week (mean weekly exercise 428.5 minutes ± 215.6; mean ± SD, range = 150 – 900 minutes), consisting primarily of sports/physical activities known to increase CRF level (e.g. running, swimming, and cycling). The low PA group comprised individuals self-reporting less than 60 minutes of deliberate exercise each week (mean weekly PA 13.8 minutes ± 25.0, range = 0 – 60 minutes; Mann-Whitney test, p = 3×10^-8^). All participants were required to complete the long format of the International Physical Activity Questionnaire (IPAQ) (Craig et al., 2003) to verify self-reported PA prior to participation.

### Experimental design

A within-subject cross-over design was used to investigate the effect of multi-day rTMS on WM. Participants completed two rTMS conditions over the course of two separate weeks, targeting the left lateral parietal cortex in one week and pre-SMA in the other week. The order of these conditions was counterbalanced across subjects and each condition was separated by an interval of at least one week (mean = 1.78, range 1 - 5 weeks). To investigate whether multi-day rTMS induced changes to local GABA concentration, MR spectroscopy assessments were conducted during the first week. Thus, changes to local GABA concentration were analysed using a between-subjects design (see Figure 1).

**Figure 1.**
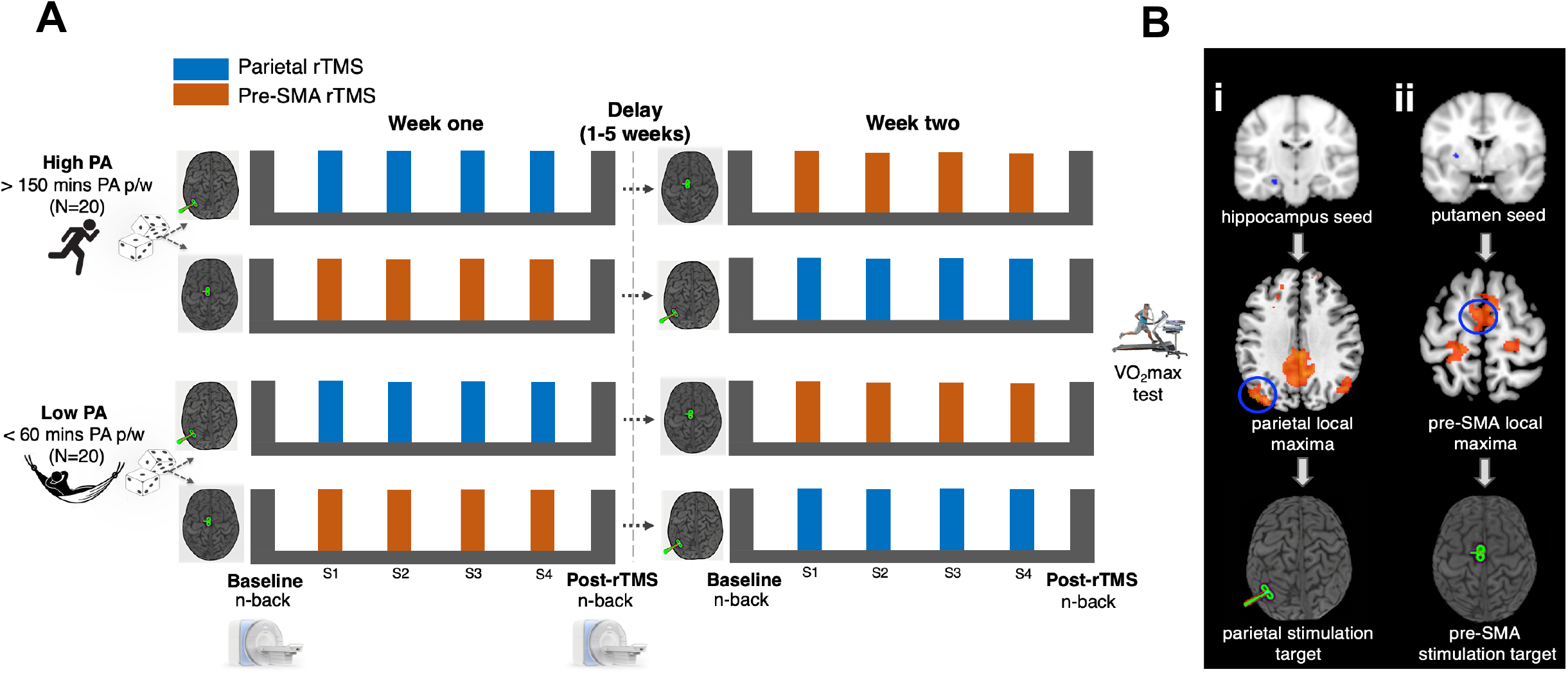
Experimental overview. **A) Multi-day rTMS protocol.** Each participant completed two multi-day rTMS conditions (S1-S4) across two separate weeks. Stimulation was applied to left lateral parietal cortex in one week, and pre-SMA in the other week. The order of these conditions was counterbalanced and pseudo-randomised within and between physical activity groups (High PA, Low PA). A minimum one-week break separated the two conditions. Working memory performance (n-back) was assessed at baseline (∼1 hour prior to S1) and post-rTMS (∼24 hours following S4). MRI assessments of GABA+ concentration were completed in week one at baseline (∼72 hours prior to S1) and post-rTMS (∼24 hours after S4). **B) rTMS target localisation.** Cortical stimulation sites were determined for each subject on the basis of individual resting-state functional MRI connectivity maps. Parietal stimulation sites were derived from a cortico-hippocampal network **(i)** and pre-SMA stimulation sites were derived from a functionally and spatially distinct cortico-striatal pre-motor network **(ii)**.

Participants were pseudo-randomised into parietal and pre-SMA stimulation conditions using an online algorithm (www.rando.la). The algorithm pseudo-randomised participants to minimise variance between conditions with regard to sex, age, level of education, preferred time of day for rTMS administration (i.e. afternoon or morning), and self-reported physical activity level (IPAQ). Furthermore, each participant’s PA categorisation (i.e. high PA vs low PA) was included in the pseudo-randomisation to achieve equal samples of high and low PA participants between week one stimulation conditions. Each condition included daily rTMS sessions for four consecutive days (i.e. Monday – Thursday), with assessments of WM at baseline and post rTMS. Baseline assessments were conducted ∼1 hour prior to the first rTMS session (i.e. Monday) and post rTMS assessments were conducted on Friday, one day after the final rTMS session (parietal condition mean delay = 24.8 ± 2.73 hrs, (mean ± SD; range = 16.5 - 29); pre-SMA condition mean delay = 24.7 ± 2.74 hrs, (mean ± SD; range = 18.5 – 29.5; p = .71).

MR-spectroscopy scans were conducted at baseline and post-rTMS to derive estimates of GABA concentration (first stimulation site only). Baseline scans were conducted on Fridays whenever possible i.e. three days prior to the first rTMS session (interval between baseline MRI and first rTMS session = 75.38 ± 11.03 hrs, [mean ± SD]; range = 66.5 - 148) and post-rTMS scans one day after the final rTMS session (interval between post MRI and final rTMS session = 23.2 ± 3.21 hrs [mean ± SD]; range = 18 - 30). Notably, previous studies have demonstrated the reliability/stability of MRS-spectroscopy GABA estimates across different timepoints (i.e. at different times of day (Evans, McGonigle, & Edden, 2010), weeks (Greenhouse, Noah, Maddock, & Ivry, 2016; Henry, Lauriat, Shanahan, Renshaw, & Jensen, 2011), and months (Near, Ho, Sandberg, Kumaragamage, & Blicher, 2014)) in both frontal (Evans et al., 2010; Greenhouse et al., 2016) and posterior brain regions (Evans et al., 2010; Greenhouse et al., 2016; Near et al., 2014). At the end of the experimental protocol, participants completed a VO_2_max test to quantify their CRF level.

### Working memory task – 2-back

To assess changes to WM following multi-day rTMS, we used an n-back task (developed in PsychoPy2 v1.85; Peirce, 2007). Participants were presented with a sequence of letters in white font in the centre of the screen on a black background. Each letter was presented individually for 1 second, with a 0.5 second inter-trial interval. Subjects were required to press the down arrow on a keyboard as quickly as possible when the letter matched a letter presented two trials previous (2-back condition). The task consisted of 130 trials containing 32 target stimuli. Alternative versions of the task with different letter sequences were used for baseline and post-rTMS assessments for each stimulation condition (and for a follow-up assessment ∼1.5 weeks following the second condition). The order of each set was randomised across subjects using a Latin square. To familiarise subjects with the task and minimise practice effects, a shortened version consisting of 33 trials containing 8 target stimuli was completed prior to the week-one baseline assessment.

### Magnetic resonance imaging (MRI)

MRI data were collected from a Siemens 3T Skyra scanner with 32-channel head coil. For both baseline and post rTMS scans, GABA-edited Mescher-Garwood Point Resolved Spectroscopy (MEGA-PRESS) data were acquired from 2.5 x 2 x 2cm voxels localised to a cortical area within left parietal cortex and pre-SMA (i.e. cortical regions encompassing rTMS targets). Voxel locations were selected on the basis of anatomical landmarks, as determination of rTMS stimulation targets required offline processing of individual resting-state fMRI data (see figure 2). The left parietal voxel was positioned in line with the lateral sulcus in the sagittal projection. The most posterior aspect of the pre-SMA voxel was positioned perpendicular to the anterior commissure, with equal coverage over left and right hemispheres (see figure 2, panel 1). 96 ON-OFF averages were acquired from each voxel position separately with TE = 68 ms, TR = 1.5 s, and an edit pulse bandwidth = 45 Hz. Editing pulses were applied at 1.9 ppm and 7.5 ppm. Following each acquisition, 8 ON-OFF averages with unsuppressed water were also obtained using the same parameters.

**Figure 2.**
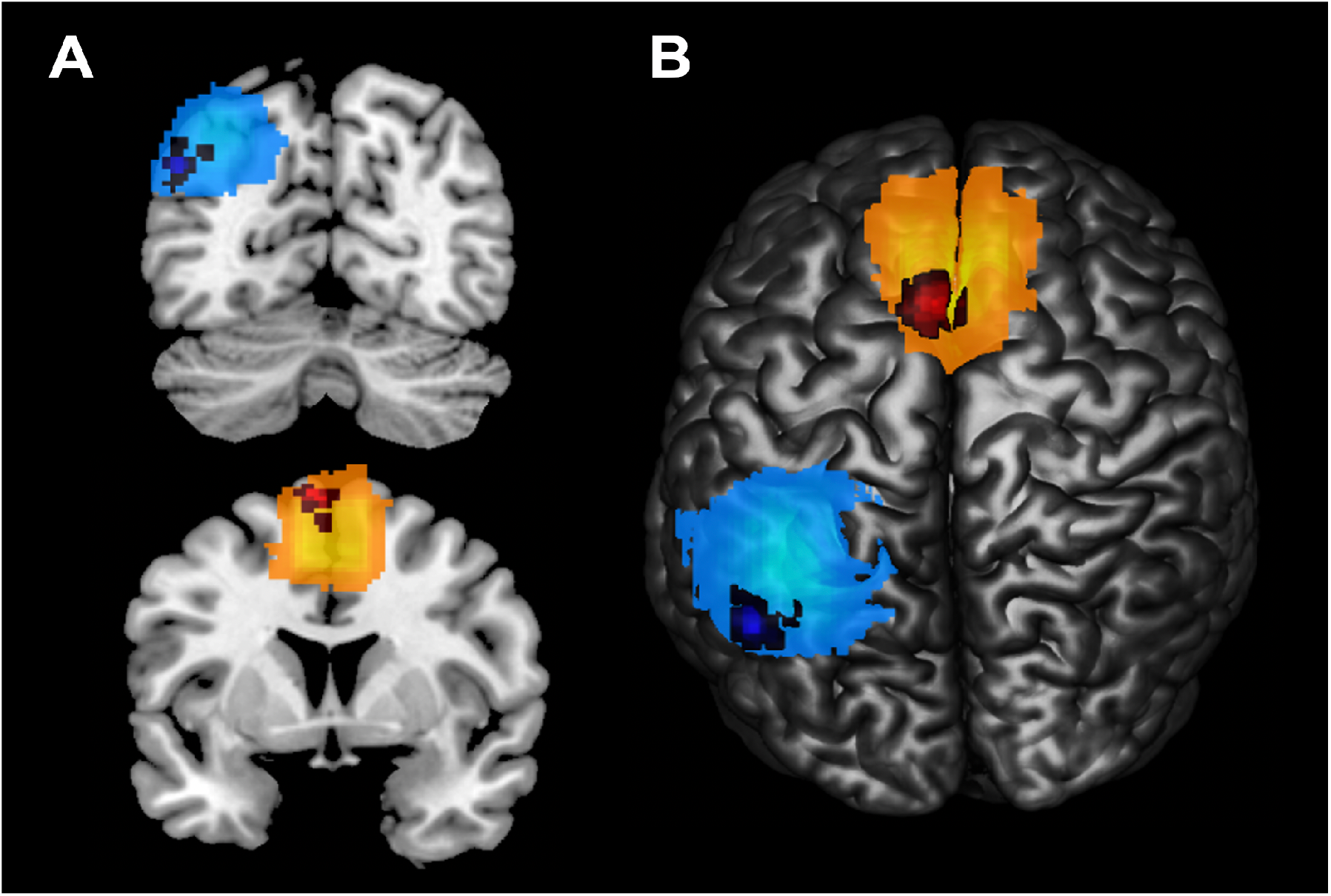
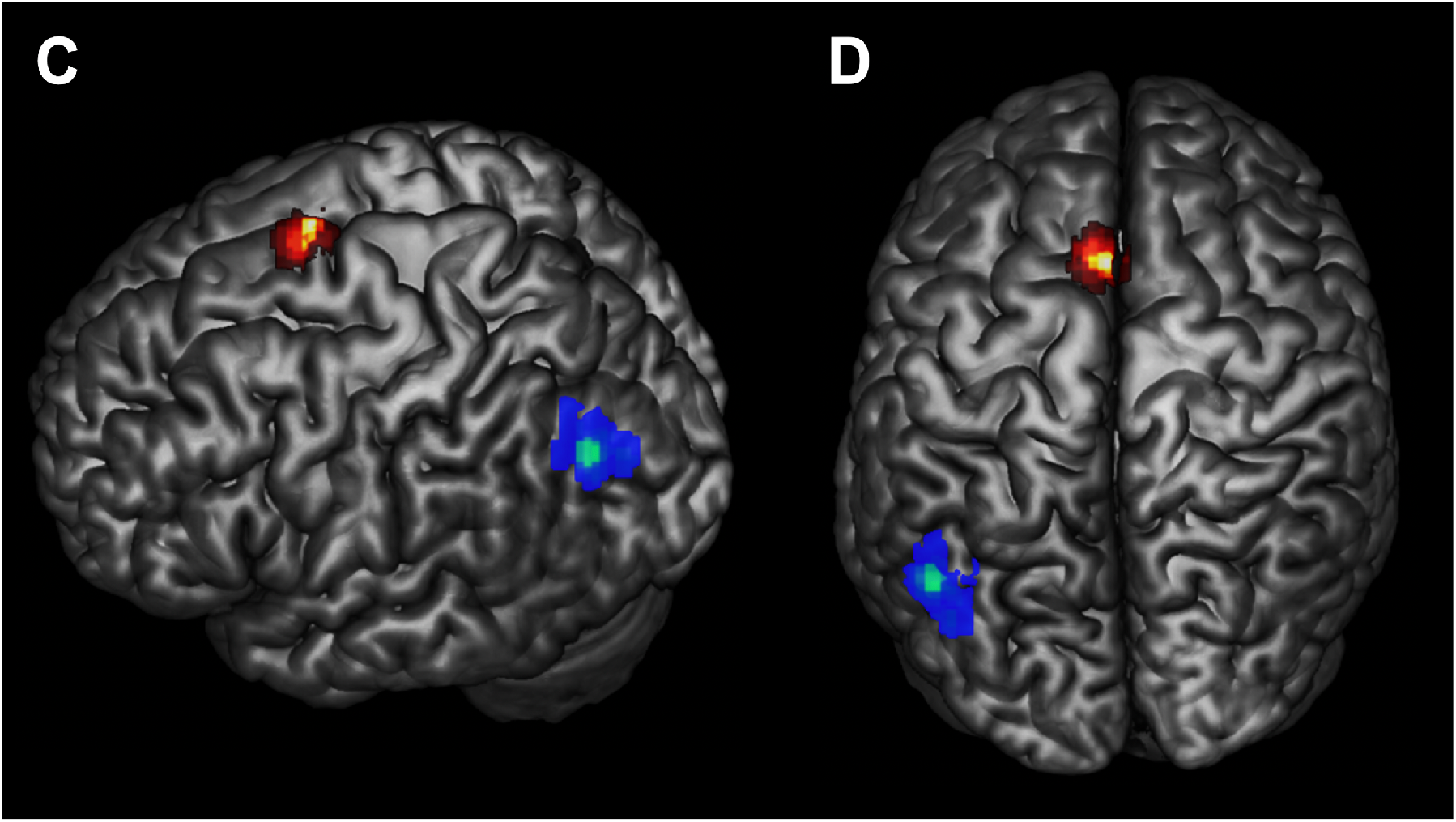
Overlay of MRS voxel positions and individualised stimulation targets. **A and B) Correspondence between MRS voxel and individualised stimulation targets.** During week one, GABA-edited MEGA-PRESS data were acquired at baseline and post rTMS from 2.5 x 2 x 2cm voxels localised to left parietal cortex and pre-SMA. The overlap in MRS voxel positions is shown (parietal voxel displayed in blue/light blue, pre-SMA displayed in orange/yellow), with greater proportion of overlap represented by lighter colouration. Stimulation targets for week one are superimposed on the corresponding MRS voxel. **C and D) Overlay of individual stimulation targets for the cross-over design.** Parietal stimulation targets were selected within a 15mm radius of MNI coordinate x = -47, y = -68, z = 36. Voxels in blue to green represent the average spatial overlap of the parietal target, with greater proportion of overlap represented in green. Pre-SMA targets were localised to an area on the cortical surface within left hemisphere, anterior to the anterior commissure (i.e. a positive y MNI coordinate). Voxels in red to yellow represent the average spatial overlap of the pre-SMA target, with greater proportion of overlap represented by yellow colouration.

T1-weighted structural images (Magnetization Prepared Rapid Gradient Echo, TR = 2.3 s, TE = 2.07 ms, voxel size 1mm^3^, flip angle 9°, 192 slices) and resting-state Echo Planar Images (TR = 2.46 s, TE = 30 ms, voxel size 3mm^3^, flip angle 90°, 44 slices) with whole brain coverage, were also acquired at both timepoints. T1-weighted images were acquired to assist with voxel localisation, while resting-state fMRI images were obtained to determine individualised stimulation targets on the basis of functional connectivity. During resting-state scans, subjects were instructed to keep their eyes open and focus on a black fixation cross presented on a white background, whilst trying not to think of anything in particular.

### Identification of rTMS stimulation locations using resting-state fMRI

Cortical stimulation sites were determined for each participant on the basis of individual resting-state functional MRI connectivity, calculated from baseline MRI data, as reported previously (Hendrikse et al. 2020). Parietal stimulation targets were derived from a cortico-hippocampal network (Wang et al., 2014) and pre-SMA stimulation sites were derived from a distinct cortico-striatal pre-motor network (Di Martino et al., 2008).

Parietal targets were defined as the cluster of voxels displaying peak local connectivity within a 15mm radius of MNI coordinate x = -47, y = -68, z = 36 (Wang et al., 2014). Across subjects the average (±SD) MNI coordinate for parietal stimulation was x = -46.4 (2.9), y = - 69.8 (3.7), z = 36.3 (3.8). Pre-SMA targets were localised to an area on the cortical surface within left hemisphere, anterior to the anterior commissure (i.e. a positive y MNI coordinate), displaying peak connectivity to the putamen seed. Pre-SMA stimulation was biased towards left hemisphere to ensure consistency between the two stimulation conditions (i.e. ipsilateral relationship between sub-cortical seed and stimulation site). Across participants the average (±SD) MNI coordinate for pre-SMA stimulation was x = -4.6 (2.7), y = 2.9 (2.4), z = 69.7 (4.1), conferring with past studies that have localised pre-SMA to 1-4cm anterior to the scalp vertex (Hamada, Ugawa, & Tsuji, 2009; Matsunaga et al., 2005) (see figure 2, panel 2).

### Repetitive transcranial magnetic stimulation

Each participant’s resting-motor threshold was determined at the beginning of each session using electromyography recorded from the right first dorsal interosseous (FDI) muscle. Biphasic pulses were applied to the left primary motor cortex (M1) using a MagVenture MagPro X100 stimulator and B-65 figure-8 cooled coil (75mm outer diameter). The motor ‘hotspot’ was determined as the area of scalp that produced the largest and most consistent motor-evoked potential in the targeted FDI muscle, and resting motor threshold was defined as the minimum stimulation intensity necessary to evoke a potential with a peak-to-peak amplitude of ≥ 50 µV in the resting FDI in at least 5 out of 10 consecutive trials.

For both parietal and pre-SMA conditions, rTMS was administered at 100% of resting-motor threshold at 20 Hz (2s on, 28s off) for 20 minutes (1600 pulses total) daily for four consecutive days. These parameters were informed by a previous study reporting improvements in memory function and network connectivity following multi-day rTMS (Wang et al., 2014), and conform to internationally established safety guidelines (Rossi, Hallett, Rossini, & Pascual-Leone, 2009). EMG recordings from the FDI muscle were continuously monitored throughout rTMS for evidence of altered motor activity (i.e. kindling). A stereotactic neuronavigation system was used throughout each session to ensure accurate target localisation relative to each subject’s neuroanatomy (Brainsight, except subjects 1-9 for whom a Zebris system was used). Stimulation targets were loaded onto each participant’s normalised T1-weighted image and aligned to the middle of the gyral crown to maximise the field size induced in cortical grey matter by the TMS pulse (Thielscher, Opitz, & Windhoff, 2011). For the parietal condition, stimulation was delivered to the target site with the coil handle perpendicular to the long axis of the gyrus to induce posterior/anterior current flow. For the pre-SMA condition, the coil was held with the coil handle pointing left, i.e. perpendicular to the midsagittal plane.

### Cardiorespiratory fitness assessment (VO_2_max)

To determine each participant’s CRF level, maximal oxygen consumption (VO_2_max) was assessed by a graded maximal exercise test conducted on a motorised treadmill. VO_2_max was conducted to provide an objective measure of individual fitness to correlate with WM and GABA outcomes. In regard to the graded exercise protocol, the starting pace was set at 6km/h with a 1% gradient, and workload was increased every three minutes (speed increase of 2 km/h until 16 km/h, followed by incremental gradient increase of 2.5%) until volitional exhaustion was reached. Heart rate and expired volume and concentration of oxygen and carbon dioxide were continuously recorded throughout the test duration (Powerlab, ADI Instruments, Dunedin, NZ). For determination of VO_2_max on the basis of VO_2_peak, at least two of the following indicators needed to be observed: (1) a plateau in VO_2_ml/kg/min values irrespective of increased workload, (2) maximal respiratory exchange ratio of ≥ 1.1, (3) a heart rate within 10 beats of age-predicted heart rate max (Tanaka, Monahan, & Seals, 2001), or (4) a self-reported rating of perceived exertion (BORG scale; Borg, 1982) rating of perceived exertion > 17 out of 20. Three subjects from the low PA group did not meet these criteria and recorded a submaximal test outcome. For these participants, individual regression equations were used (Mean R^2^ = .77) to derive an estimated VO_2_max value on the basis of age-predicted max heart rate (Tanaka et al., 2001).

### Data analysis

#### Cardiorespiratory fitness

Two subjects from the high PA group were lost to follow-up and did not complete the graded exercise test. Thus, analyses were conducted with N = 38 (high PA, n = 18; low PA, n = 20). To assess differences in fitness level between low PA and high PA groups, independent-samples t-tests were conducted on i) VO_2_max scores and ii) total test duration (mins). We also assessed the association between VO_2_max and current self-reported weekly exercise using a Pearson’s correlation.

#### Working memory task - 2-back

Performance on the task was scored by reaction time to correct responses (hits), and *d’* values (z-transformed percentage of hits relative to z-transformed percentage of false alarms, *d’* = Z_hits_ – Z_false alarm_). For one subject, scheduling conflicts meant that it was not feasible to conduct post-rTMS assessments 24 hours following the final rTMS session. Data from another subject was removed following misinterpretation of task instructions and an erroneous response pattern (i.e. responding to each trial, resulting in inflated false alarm rates). Thus, data analyses of WM change were conducted on N = 38.

We compared reaction time (correct trials) and *d’* values across stimulation conditions in the cross-over design in a 2 x 2 x 2 repeated-measures ANOVA, with within-subject factors of stimulation condition (parietal, pre-SMA), time (baseline, post-rTMS), and a between-subject factor of physical activity (high PA, low PA).

To account for the possibility of carry-over effects between parietal and pre-SMA stimulation conditions, we also assessed changes to WM following the first stimulation condition. Specifically, baseline versus post-rTMS assessments were compared between stimulation and physical activity groups in a 2 x 2 x 2 repeated measures ANOVA, with the within-subject factor of time (baseline, post-rTMS) and between-subject factors of stimulation condition (parietal, pre-SMA) and physical activity (high PA, low PA). Further, to assess whether stimulation induced lasting effects, independent-measures ANOVAs were conducted on follow-up assessments of reaction time and *d’* (∼1.5 weeks following final rTMS session), with between-subject factors of stimulation condition (parietal, pre-SMA) and physical activity (high PA, low PA).

#### GABA quantification

Gannet (version 3) (Edden et al., 2014) was used to analyse MEGA-PRESS spectra. Estimates of GABA concentration (3.0 ppm) were calculated for each voxel location (left parietal cortex and pre-SMA), relative to the unsuppressed water signal. The Gannet package includes two modules, GannetLoad and GannetFit, which conduct pre-processing and quantification of GABA spectra respectively. GannetLoad module was employed for phased-array channel combination of raw Siemens ‘twix’ files, exponential line broadening (3 Hz), Fourier transformation to yield time-resolved frequency-domain spectra, frequency and phase correction, outlier rejection, and time averaging of the edited difference spectrum. GannetFit module was employed to estimate the area of the edited GABA peak at 3 ppm using a single Gaussian peak model, and a nonlinear least-squares fitting approach. To acknowledge a possible macromolecule contribution to the edited spectra, GABA+ is referred to in place of GABA for all relevant analyses (Edden, Puts, Harris, Barker, & Evans, 2014). As an internal reference, GABA+ concentrations are expressed in institutional units, relative to the unsuppressed water signal, which was estimated with a Gaussian-Lorentzian model.

Partial volume segmentation of the T1-weighted anatomical image within each voxel was conducted using FSL’s FAST(Jenkinson, Beckmann, Behrens, Woolrich, & Smith, 2012), and FSLmaths was used to calculate the percentage of voxel overlap between baseline and post-rTMS assessments for parietal and pre-SMA voxels. GABA+:H_2_O ratios were partial volume corrected by removing cerebrospinal fluid fractions from GABA+ estimates (GABA+:H_2_O / (1 - CSF%)). The full width at half maximum of GABA signals (FWHM) and GABA signal fit error (relative to the unsuppressed water peak, E_GABA_) were used to assess the quality of GABA+ estimates. Spectra with an E_GABA_ value of > 20% or exceeding a z-score of 3.29 were labelled as outliers and removed from the analysis. To assess stability of GABA+ estimates across time and between stimulation conditions, 2 x 2 x 2 x 2 repeated measures ANOVAs were conducted on data quality metrics (FWHM, E_GABA_), with between-subject factors of Stimulation Condition (parietal, pre-SMA), Physical Activity (high PA, low PA), and within-subject factor of Time (baseline, post-rTMS) and Voxel Location (parietal, pre-SMA). We also assessed the stability of the water reference signal. Water FWHM values were examined across Stimulation Conditions, Time, and Voxel Sites in the same 2 x 2 x 2 x 2 repeated-measures ANOVA model.

To investigate the effects of multi-day rTMS on GABA+ estimates, a 2 x 2 x 2 x 2 repeated measures ANOVA was conducted with within-subject factors of Time (baseline, post-rTMS) and Voxel Location (parietal, pre-SMA), and between-subject factors of Stimulation Condition (parietal, pre-SMA) and Physical Activity (high PA, low PA). To account for differences in baseline GABA+ concentration between individuals in the different stimulation conditions, we applied a separate analysis where difference scores were calculated by subtracting baseline GABA+ estimates from post-rTMS estimates (i.e post-rTMS – baseline). Differences scores were entered into a 2 x 2 x 2 repeated-measures ANOVA, within-subject factor of Voxel Site (parietal, pre-SMA) and between-subject factors of Stimulation Condition (parietal, pre-SMA) and Physical Activity (high PA, low PA).

#### Association between working memory, GABA+ and cardiorespiratory fitness

We assessed if changes (post-rTMS – baseline) to WM and GABA+ concentration following multi-day rTMS were related to individual CRF level (VO_2_max) using Pearson’s correlations. GABA+ estimates were taken from the voxel site corresponding to the stimulation target (i.e. parietal voxel for subjects who received parietal stimulation, and pre-SMA voxel for subjects who received pre-SMA stimulation). Further, Pearson’s correlations were also conducted to determine whether changes to GABA+ were associated with improved WM.

Statistical analysis of these data was conducted using JASP (v 0.10.2). An alpha level of .05 was adopted for all inferential statistics conducted under the null hypothesis significance testing framework. To quantify the relative evidence for the alternative vs. the null hypothesis, equivalent Bayesian statistics were also conducted (prior set in support of the alternative over null hypothesis, i.e. BF_10_). For Bayesian tests, BF_10_ > 3 was considered strong evidence against the null, and BF_10_ < 0.33 strong evidence for the null. For Bayesian t-tests, the Cauchy parameter was set to a conservative default value of 0.707 (Ly, Verhagen, & Wagenmakers, 2016; Rouder, Speckman, Sun, Morey, & Iverson, 2009).

## Results

Data were collected from a sample of 40 participants (19 males, 21 females) with an average age of 25.48 (SD ± 9.35) years. Both rTMS conditions were well tolerated by subjects; two subjects reported transient mild headaches following stimulation (one following parietal rTMS, and one following pre-SMA rTMS), but no other adverse events were reported.

Due to scheduling conflicts, 2/40 subjects did not complete the MRI protocol. Therefore, MRI analyses were conducted with N = 38, with comparisons between individuals who received parietal stimulation (n = 19) versus pre-SMA stimulation (n = 19). There were no significant differences between groups in regard to age or sex (see table 1 in supplementary materials).

**Table 1:** average TMS intensities. Percentage of maximum stimulator output (MSO) (MagVenture X100, Biphasic) at which rTMS was administered (shown for within-subject comparisons, week one stimulation conditions, and PA groups). Parentheses indicate standard deviation.

|  | TMS intensity (% MSO) |
| --- | --- |
| Within-subjects cross-over (N = 38) | 47.24% (8.16) |
| Parietal stimulation Week 1 (N = 19) | 46.89% (8.26) |
| Pre-SMA stimulation Week 1 (N = 19) | 47.58% (8.26) |
| High PA (N = 19) | 46.11% (6.24) |
| Low PA (N = 19) | 48.37% (9.75) |

TMS intensities (% of maximum stimulator output) for the complete cross-over sample, week one stimulation conditions, and PA groups are reported in table 1. There was no significant difference in TMS intensity between week one parietal and pre-SMA stimulation groups (t_1,36_ = -0.26, p = .80, Cohen’s d = 0.08), nor between high PA and low PA groups (t_1,36_ = -0.85, p = .40, Cohen’s d = 0.28).

### Differences in cardiorespiratory fitness level between physical activity conditions

We verified that CRF differed between high PA and low PA groups with independent t-tests. VO_2_max and graded exercise test duration were greater in the high PA group relative to the low PA group (see table 2). Further, there was a significant positive association between self-reported current weekly exercise and VO_2_max, with increasing weekly exercise minutes associated with greater CRF (r = .61, p = 4.29 x 10^-5^) (figure S1).

**Table 2:** Comparison of cardiorespiratory fitness between High and Low PA groups

| Outcome Measure | Mean (SD)<br>High PA | Mean (SD)<br>Low PA | Statistic | df | p | Effect size | BF<sub>10</sub> |
| --- | --- | --- | --- | --- | --- | --- | --- |
| VO<sub>2</sub>max<br>(ml/kg/min) | 52.11(9.11) | 38.26 (9.06) | 4.70<sup>a</sup> | 1,36 | **3.80 x 10<sup>-5</sup>** | 1.53<sup>d</sup> | **496.05** |
| Test<br>Duration<br>(mins) | 15.14 (2.90) | 9.55 (2.82) | 6.02<sup>a</sup> | 1,36 | **6.53 x 10<sup>-7</sup>** | 1.96<sup>d</sup> | **18218.50** |

<sup>a</sup> *t* statistic (Independent-samples *t*-test); <sup>b</sup> Cohen's *d*; Time VO<sub>2</sub> = total graded exercise test duration (mins); bolded values indicate *p* < .05

### Working memory – n-back

We assessed correct trial reaction times and *d’* across stimulation conditions in the cross-over design (i.e. within-subjects, parietal versus pre-SMA stimulation) with 2 x 2 x 2 repeated-measures ANOVAs. There were no main effects or interactions for either reaction time or *d’* scores (see table 3, figure 3). To account for the possibility of carry-over effects between stimulation conditions, we also assessed working memory following the first stimulation condition only (i.e. between-subject factors of stimulation site, and physical activity group). There were significant interactions between Stimulation Condition and Time for both reaction time and *d’ (*p < .05, BF_10_ > 1.60), with post-hoc tests indicating that this was driven by an increase following pre-SMA stimulation (see supplementary materials). However, these effects were not seen at follow-up ∼15 days following the final stimulation session (figure S3).

**Figure 3.**
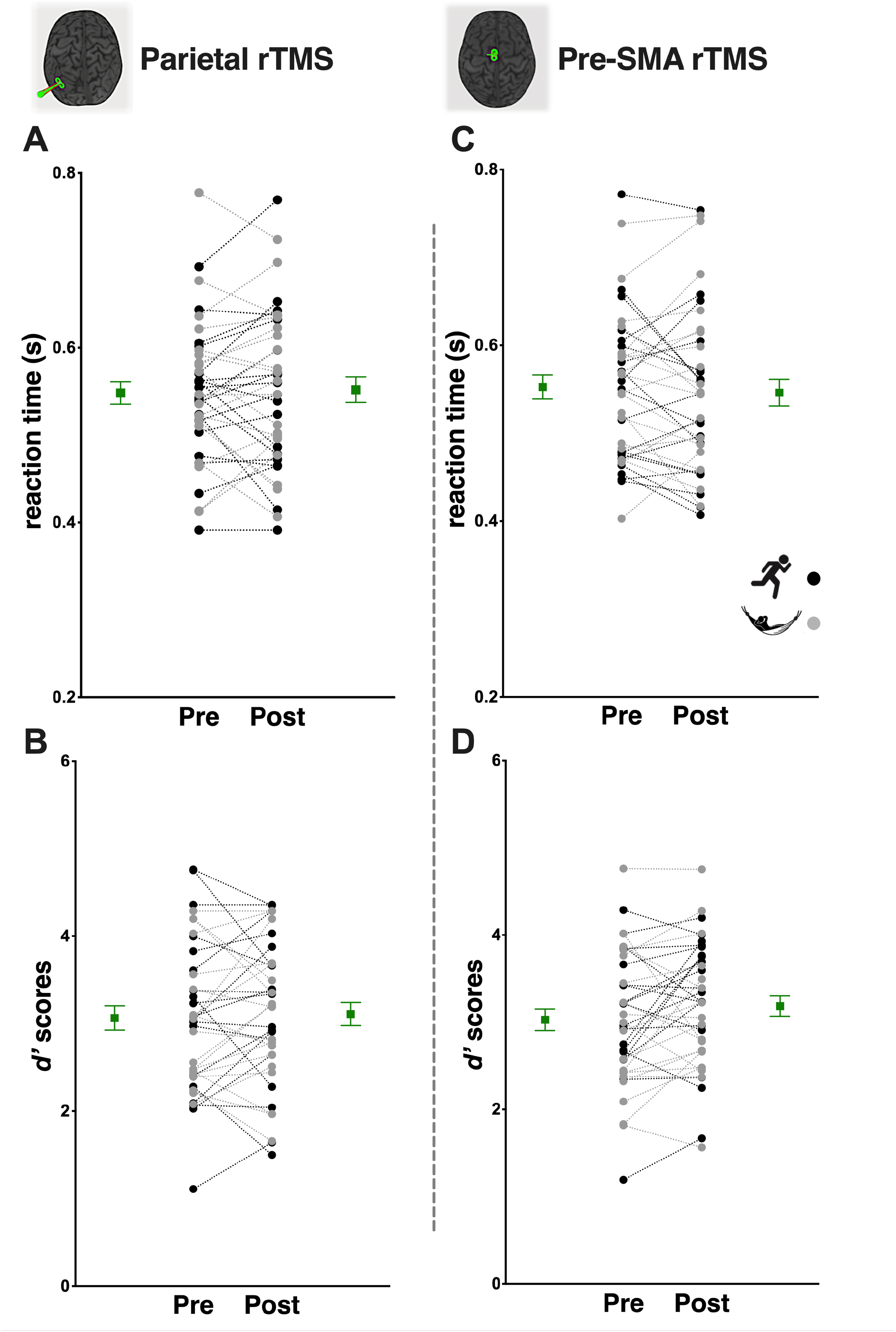
Effects of multi-day rTMS and physical activity on working memory. No evidence of improved working memory function following multi-day rTMS when comparing across stimulation conditions in the cross-over design. The left panel **(A,B)** shows performance on the n-back task following parietal rTMS (**A,** reaction time; **B,** *d’* scores). The right panel **(C,D)** shows performance on the n-back task following pre-SMA rTMS (**C,** reaction time; **D,** *d’* scores). Circles represent individual scores with high PA individuals depicted in black and low PA individuals in grey; squares and error bars depict group mean and standard error.

**Table 3:** Working memory performance (reaction time, *d’*) in cross-over design

| Cases | F ratio | df | p | $\eta^2_p$ |
| --- | --- | --- | --- | --- |
| *Reaction time* |  |  |  |  |
| Time | 0.05 | 1,36 | .82 | .00 |
| Stim. Cond. | 0.01 | 1,36 | .92 | .00 |
| PA | 0.33 | 1,36 | .57 | .01 |
| Time x Stim. Cond. | 0.82 | 1,36 | .37 | .02 |
| Time x Stim. Cond. x PA | 1.20 | 1,36 | .28 | .03 |
| *$d'$* |  |  |  |  |
| Time | 2.69 | 1,36 | .11 | .07 |
| Stim. Cond. | 0.06 | 1,36 | .81 | .00 |
| PA | 0.49 | 1,36 | .49 | .01 |
| Time x Stim. Cond. | 0.82 | 1,36 | .37 | .02 |
| Time x Stim. Cond. x PA | 0.46 | 1,36 | .50 | .01 |

Stim. Cond. = Stimulation Condition; PA = Physical Activity

**Table 3:** The effects of multi-day rTMS on GABA+ concentration

| Cases | F ratio | df | p | $\eta^2_p$ | BF<sub>10</sub> |
| --- | --- | --- | --- | --- | --- |
| Time | 2.07 | 1,31 | .16 | .06 | - |
| Stim. Cond. | 0.74 | 1,31 | .40 | .02 | - |
| PA | 0.19 | 1,31 | .66 | .01 | - |
| Voxel Site | 4.55 | 1,31 | **.041** | .13 | 1.73 |
| Time x Stim. Cond. | 5.26 | 1,31 | **.029** | .15 | 1.76 |
| Time x Stim. Cond. x Voxel Site | 0.96 | 1,31 | .34 | .03 | - |
| Time x Stim. Cond. x PA | 4.33 | 1,31 | **.046** | .12 | 1.27 |
| Time x Stim. Cond. x Voxel Site x PA | 4.65 | 1,31 | **.039** | .13 | 2.21 |

Stim. Cond. = Stimulation Condition; PA = Physical Activity; bolded values indicate $p < .05$

In summary, our findings suggest that multi-day rTMS to the parietal cortex did not improve WM performance. However, there was some evidence of improved WM following multi-day rTMS to pre-SMA, although improvements were only observed following the first week of stimulation, and not in the full cross-over design.

### Effects of stimulation and physical activity on GABA+ concentration

We first assessed MRS data quality. Voxel locations were consistent across time points, and stimulation and physical activity groups (p > .10 for all main effects/interaction terms). There were also no significant differences in tissue fractions (grey matter, white matter, and CSF) across time points and stimulation conditions (p > .42 for all main effects/interaction terms; see supplementary materials for all data quality analyses). Regarding GABA+ concentrations, three outlier values were identified (values outside the physiologically plausible range, z-scores > +3.29) for the pre-SMA voxel (1 from the baseline timepoint, 2 from post-rTMS time point); these values were removed from all analyses. To assess the effects of multi-day rTMS and physical activity on GABA+ concentration, a 2 x 2 x 2 x 2 repeated-measures ANOVA was conducted (see table 3).

Post-hoc tests revealed higher GABA+ concentrations in the pre-SMA voxel relative to the parietal voxel (Cohen’s d = 0.36; p = .041 [Tukey corrected post hoc test], BF_10_ = 1.73), and a trend towards a generalised increase in GABA+ across voxels following pre-SMA stimulation (Cohen’s d = 0.45; p = .054 [Tukey corrected post hoc test], BF_10_ = 1.76). No other significant differences were observed between levels for other interaction effects (Tukey’s tests, all p > .09) (see figure 5).

**Figure 5.**
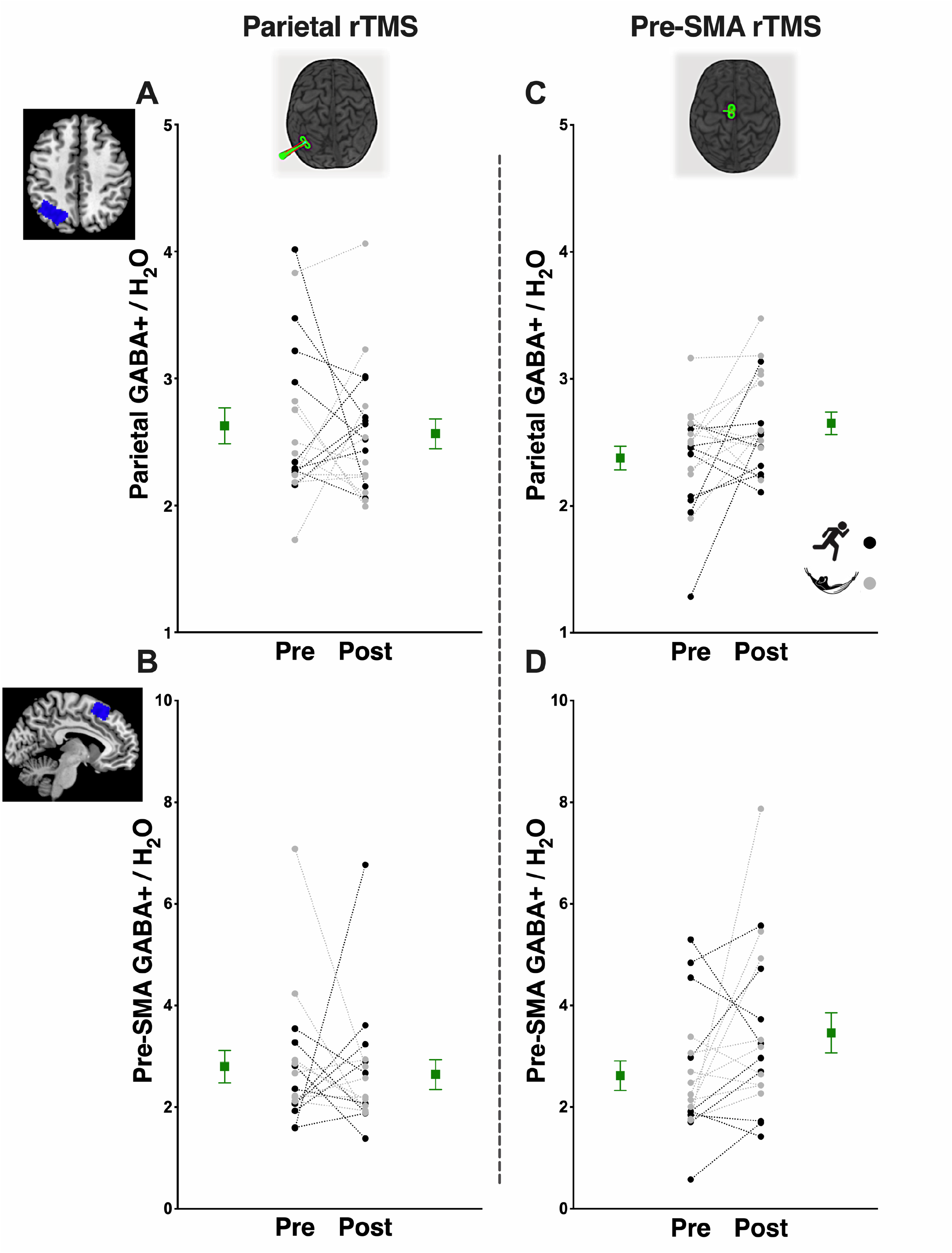
Effects of multi-day rTMS and physical activity on GABA+ concentrations. No evidence of site-specific effects of multi-day rTMS on GABA+ concentration. However, a trend towards a generalised increase in GABA+ following multi-day pre-SMA rTMS. The left panel **(A,B)** shows GABA+ estimates following parietal rTMS (**A**, parietal voxel; **B**, pre-SMA voxel). The right panel **(C,D)** shows GABA+ estimates following pre-SMA rTMS (**C**, parietal voxel; **D**, pre-SMA voxel). Circles represent individual scores with high PA individuals depicted in black and low PA individuals in grey; squares and error bars depict group mean and standard error.

To account for variability in baseline estimates between stimulation conditions, we also conducted a 2 x 2 x 2 repeated-measures ANOVA on each subject’s change in GABA+ concentration at each voxel location (post-rTMS GABA – baseline GABA). We observed a significant difference in Δ pre-SMA GABA+ concentration between stimulation conditions, with evidence of an increase in pre-SMA GABA+ concentration following pre-SMA stimulation in low PA individuals (p = .011, Cohen’s d = 0.62, BF_10_ = 4.01) (figure S4).

Overall, our findings suggest that multi-day rTMS did not exert site-specific effects on GABA+ concentration. In contrast, we found trend-level evidence of a generalised increase in GABA+ concentration across both voxel locations following multi-day rTMS to pre-SMA. Additionally, our results suggest that high levels of exercise did not enhance the effects of multi-day rTMS on GABA+ concentrations.

### Associations between working memory, GABA+, and cardiorespiratory fitness

We first examined associations across individuals who had received multi-day parietal stimulation (Figure 6). In contrast to our hypothesis, VO_2_max was not associated with change in working memory following parietal stimulation (VO_2_max vs reaction time, r = -.03, p = .88, n =36; VO_2_max vs *d’* values, r = .04, p = .36, n = 36), and VO_2_max was not associated with change in GABA+ concentration at the stimulation site (i.e. within parietal voxel) (r = .24, p = .35, n = 17). However, we did observe weak evidence for a negative correlation between changes in reaction time and GABA+ within the parietal voxel following stimulation (r = -.49, p = .045, n = 17, BF_10_ = 1.93), though this effect is non-significant when correcting for multiple comparisons (i.e. Bonferroni-adjusted α-level = .01). Thus, there is some weak evidence that greater increases in GABA+ following parietal stimulation were associated with faster response times on the working memory task.

**Figure 6.**
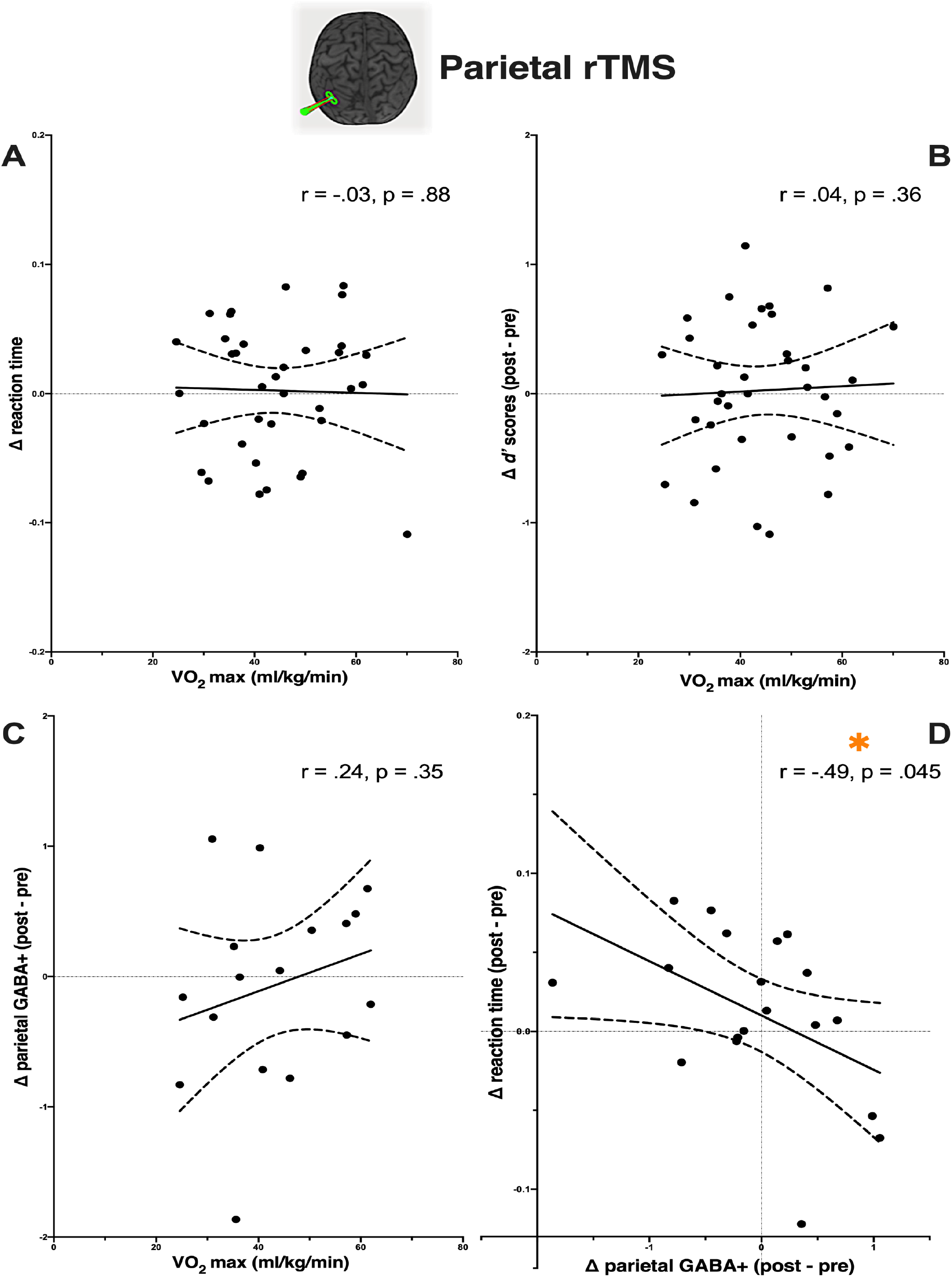
Associations between cardiorespiratory fitness level (VO_2_max) and change in working memory and GABA+ concentration following multi-day parietal rTMS. D) Improved reaction time on the n-back task was associated with greater decrease in parietal GABA following parietal stimulation. Solid line shows regression line of best fit, dotted lines depict 95% confidence interval. * p < .05.

We next examined associations within individuals who had received multi-day pre-SMA stimulation (Figure 7). There was an association between VO_2_max and change in WM following pre-SMA stimulation (VO_2_max vs reaction time, r = -.44, p = .008, n =36, BF_10_ = 6.35, Figure 7A; VO_2_max vs *d’* values, r = .36, p = .032, n = 36, BF_10_ = 1.88, Figure 7B). The association between VO_2_max and reaction time remained significant when correcting for multiple comparisons, though the association between VO_2_max and *d’* values did not (Bonferroni-adjusted α-level = .02). VO_2_max was not associated with change in GABA+ concentration at the stimulation site (i.e. within pre-SMA voxel) (r = -.25, p = .31, n = 18).

**Figure 7.**
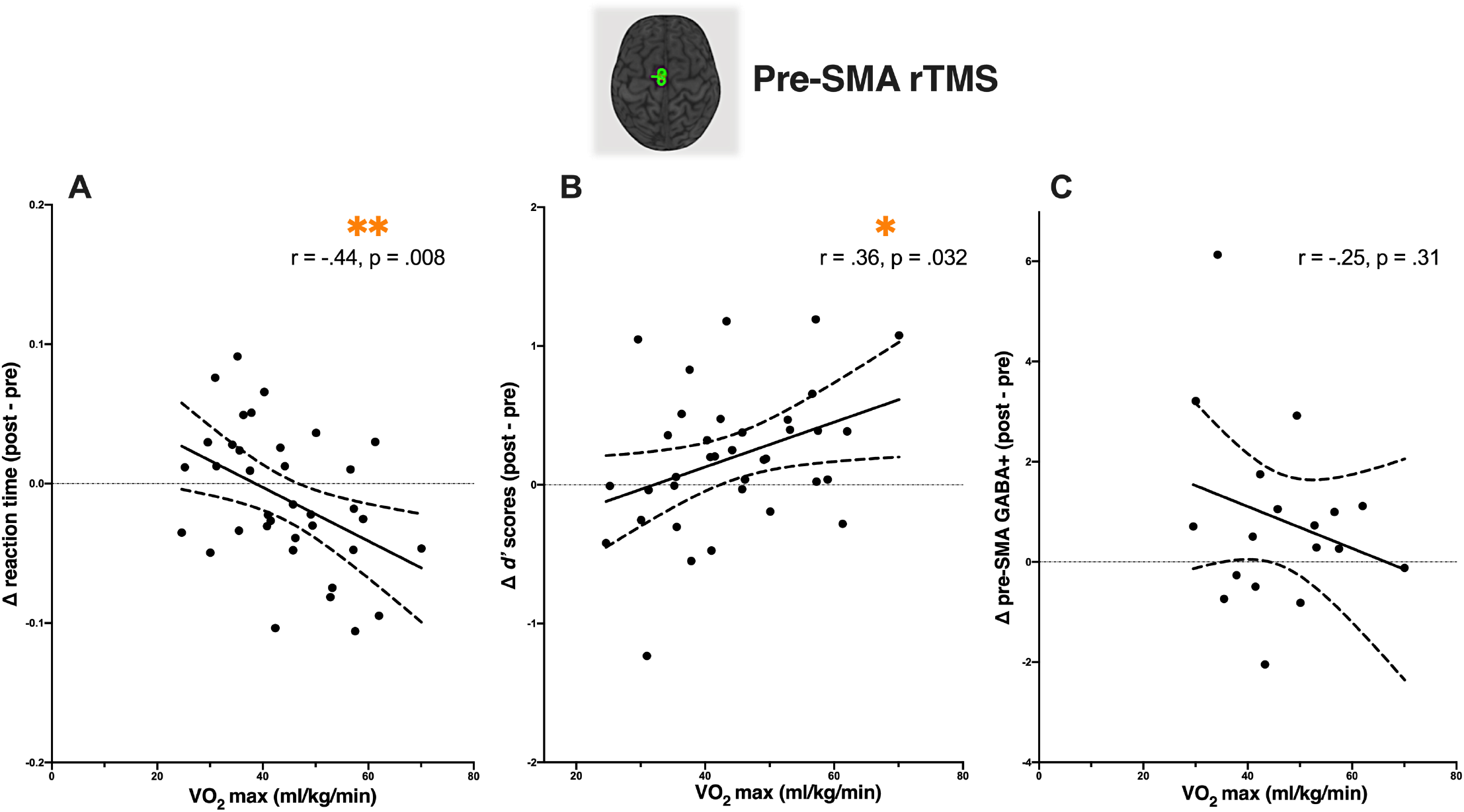
Associations between cardiorespiratory fitness level (VO_2_max) and change in working memory and GABA+ concentration following multi-day pre-SMA rTMS. Evidence of associations between individual fitness level and change in working memory **(A,B)** but not GABA+ concentration **(C)** following pre-SMA rTMS. The solid line shows regression line of best fit, dotted lines depict 95% confidence interval. * p < .05; ** p < .01.

In summary, our findings provide some evidence that increases in parietal GABA+ following multi-day rTMS may be associated with improved working memory performance. Our results also provide some indication that higher CRF is associated with greater improvement in both reaction time and *d’* values following pre-SMA stimulation.

## Discussion

In this study, we hypothesised that multi-day rTMS would improve working memory performance and increase GABA concentration. We did not observe evidence of enhanced WM following multi-day parietal stimulation. There was some evidence of improved WM function following pre-SMA stimulation, though these effects were only present for the first week of stimulation, and not in the full cross-over design. Further, in contrast to our hypothesis that there would be site-specific changes in GABA concentration following multi-day rTMS, we only observed trend-level evidence of a generalised increase in GABA concentration across stimulation sites following multi-day pre-SMA stimulation. Finally, we investigated whether regular exercise engagement modulated any stimulation effects. We expected that higher levels of physical activity and CRF would enhance the effects of multi-day rTMS. We did not observe evidence of an interaction between physical activity level and changes to WM or GABA concentration following stimulation. However, we found that higher CRF was associated with greater WM improvement following pre-SMA stimulation.

### Multi-day rTMS and working memory

A number of studies have demonstrated transient improvements in WM following a single dose of rTMS to parietal cortex (Hamidi et al., 2008; Luber et al., 2007; Yamanaka et al., 2010), as well as longer term improvements following multi-day rTMS (6 sessions; Pearce et al., 2014). We did not observe evidence of improved WM performance following multi-day parietal stimulation. Our findings are in contrast to a study by Pearce et al. (2014) which reported improved spatial WM performance following multi-day parietal rTMS. There are a number of methodological differences between studies which may have contributed to these discrepant outcomes. For example, in the current study we applied 20 Hz rTMS over four consecutive days (1600 pulses per session) to subject-specific parietal locations determined on the basis of hippocampal functional connectivity (average MNI coordinate: x = -46.4, y = - 69.8, z = 36.3). In comparison, Pearce et al. (2014) applied six sessions of 5 Hz rTMS across a two-week period (i.e. rTMS on alternative days, 300 pulses per session) to the P4 electrode site on the 10-20 electroencephalography system (i.e. right hemisphere, corresponding to MNI coordinate x = 36.8, y = -74.9, z = 49.2; Okamoto et al., 2004). It is possible that differences in stimulation parameters (i.e. stimulation site, frequency, dosage, timing), or target localisation approach, may have contributed to the differing pattern of results. It is also possible that the task demands between studies (i.e. 2-back vs Cambridge spatial WM task) were different. The study by Pearce et al. (2014) was conducted on a small sample of healthy individuals (N = 10 received parietal rTMS). Given the known inter-individual variability associated with rTMS plasticity protocols (López-Alonso, Cheeran, Río-Rodríguez, & Fernández-Del-Olmo, 2014), it is also possible that the null findings of the current study, conducted with a larger sample of 40 healthy individuals using a cross-over within-subjects design, provide a more accurate estimate of the true effect size.

Our findings provide some weak evidence of improved WM following pre-SMA stimulation. We observed evidence of improved reaction time and overall performance (*d’* values) following pre-SMA stimulation in the first week, although the improvement in performance was not present in the full cross-over design. The pre-SMA is a key region within premotor networks supporting executive control (e.g. task switching) as well as distinct forms of implicit memory (motor skill learning, procedural memory) (Hikosaka et al., 1996; Hikosaka, Nakamura, Sakai, & Nakahara, 2002). Although functional neuroimaging studies typically emphasise the contribution of a fronto-partietal network, WM studies have also demonstrated the involvement of pre-motor regions, including the pre-SMA (Buchsbaum & D’Esposito, 2019; Marvel et al., 2019; Rottschy et al., 2012). In light of our findings, it is possible that this region may provide an alternative rTMS target to the parietal cortex or dorsolateral prefrontal cortex to modulate WM. However, these effects were only observed following the first week of stimulation and not in the cross-over analysis, and may therefore reflect a false positive result. Alternatively, the null result for the cross-over analysis may reflect carry-over effects. Half of the participants received parietal cortex stimulation in the first week which may have influenced/occluded the effects of pre-SMA stimulation. Future research is needed to disentangle the effects of multi-day rTMS on WM performance across these regions.

### Multi-day rTMS and GABA+ concentration

It is well established that rTMS to primary motor cortex results in changes GABAergic neurotransmission (Daskalakis et al., 2006; Stagg et al., 2009; Ziemann et al., 2008). In contrast, there have been few attempts to investigate the effects of rTMS on GABA outside of the motor system (e.g. Casula, Pellicciari, Picazio, Caltagirone, & Koch, 2016; Chung et al., 2017). Similarly, few studies have examined the potential for multi-day rTMS to induce lasting changes to GABA concentration. A single dose of theta burst stimulation (TBS) to either the primary motor cortex (continuous TBS; Stagg et al., 2009) or the left inferior parietal cortex (intermittent TBS; Vidal-Piñeiro et al., 2015) has been reported to increase GABA concentration as measured with MRS. We extend upon these single dose studies with our finding of an increase in GABA concentration following multi-day 20 Hz pre-SMA stimulation. However, in contrast to our hypothesis, we did not observe evidence of site-specific changes, but instead report trend-level evidence of a generalised increase in GABA concentration across voxel sites.

MRS provides an estimate of the overall tissue concentration of GABA (Stagg, 2014), which on a functional level may reflect extrasynaptic ‘GABAergic tone’ (Maddock & Buonocore, 2012; Stagg et al., 2011). It is therefore possible that the increase in GABA concentration observed following rTMS occurs via modulation of tonic GABA activity (Stagg et al., 2011). rTMS is thought to induce neuroplasticity via mechanisms akin to long-term potentiation/depression (Hoogendam, Ramakers, & Di Lazzaro, 2010), in which modulation of GABA is a necessary prerequisite (Castro-Alamancos, Donoghue, & Connors, 1995; Stagg, 2014). The repeated, sustained triggering of these effects via 20 Hz multi-day rTMS may induce accumulative changes to GABA concentration (remaining visible via MRS ∼24 hours following stimulation).

In contrast to the generalised increase in GABA following multi-day pre-SMA rTMS, we did not find evidence of changes following stimulation of parietal cortex. The underlying cause of this discrepancy is unclear, though there is some evidence that GABA concentration varies across different cortical regions (Veen & Shen, 2013). However, we found that baseline GABA concentrations were consistent between voxel sites (see supplementary materials), suggesting that such regional differences are unlikely to have influenced our results. Future work is required to characterise the regional specificity of rTMS-induced changes to GABA.

### Associations between exercise and changes to working memory and GABA concentration following rTMS

Exercise upregulates a number of mechanisms which promotes an environment conducive to neuroplasticity (Hendrikse et al., 2017), including altered neurotransmission (e.g. modulation of GABA; Coxon et al., 2018; Mooney et al., 2016). Hence, in this study we examined whether high levels of exercise and CRF influenced the effects of rTMS on WM and GABA. We did not observe consistent evidence of an interaction between current self-reported level of physical activity and changes to WM or GABA concentration following stimulation (though we observed some evidence of increased GABA concentration following pre-SMA rTMS in individuals engaging in low physical activity, see figure S3 in supplementary materials). However, we did observe evidence of a positive relationship between CRF level and change in WM following pre-SMA rTMS. Specifically, we found that higher CRF was associated with greater improvement in reaction time and overall performance (*d’*) following pre-SMA rTMS. The underlying cause of the inconsistent interactions between WM, GABA, physical activity, and CRF is unclear. However, this may relate to the distinction between our measures of physical activity/exercise and CRF. To some extent, increased CRF reflects physiological adaptation within the respiratory systems in response to regular long-term engagement in moderate/vigorous aerobic activity (Carrick-Ranson et al., 2014). In contrast, our measure of self-reported weekly exercise level likely reflects a more recent pattern of exercise engagement (Shephard, 2003; Sylvia, Bernstein, Hubbard, Keating, & Anderson, 2014). It is plausible that these measures may therefore have different interactions with GABA, WM, and rTMS. However, conceptually, the evidence of a positive association between CRF and change in WM aligns with past studies demonstrating an interaction between exercise and the response to single doses of rTMS (Cirillo et al., 2009; McDonnell, Buckley, Opie, Ridding, & Semmler, 2013). While speculative, the relationship between CRF and WM provides some indication that sustained engagement in exercise may improve the effects of rTMS on cognitive function.

### Limitations and future directions

There are certain methodological limitations of this study which should be acknowledged. Our experimental design featured two active stimulation conditions, whereby 20 Hz stimulation was applied to parietal cortex and pre-SMA across two separate weeks. This protocol allowed assessment of the site-specificity of rTMS-induced changes to WM and GABA concentration. However, we acknowledge that the inclusion of a dedicated sham condition would have provided the opportunity to characterise the stability of MRS GABA estimates over time, and examine the extent to which changes in WM performance may be attributable to generalised practice or placebo effects. However, we note that previous studies have established the reliability/stability of MRS GABA estimates over repeated assessments across similar timeframes (Greenhouse et al., 2016; Henry et al., 2011). Further, to familiarise subjects with the n-back task and minimise practice effects, a shortened version was completed prior to the baseline assessment. We also compared performance using a within-subjects cross-over design (N = 38), and showed no significant changes following either parietal or pre-SMA stimulation conditions, with Bayesian statistics reporting consistent evidence in favour of the null hypothesis. Hence, we can be more confident that our experimental design and analysis has provided a reliable assessment of the effects of multi-day rTMS on GABA concentration and WM performance.

We aimed to investigate the interaction between current physical activity level and the effects of multi-day rTMS on WM and GABA concentration. To address this, we recruited individuals who were engaging in high versus low amounts of exercise as part of their normal weekly routine, and assessed their response to multi-day rTMS. Thus, the exact timing of exercise relative to rTMS administration was not standardised across individuals. While higher levels of self-reported physical activity/aerobic exercise have been shown to increase the response to rTMS (Cirillo et al., 2009), it is possible that variation in the temporal relationship between exercise engagement and rTMS administration across individuals may have impacted our findings. For instance, exercise performed prior to versus after rTMS may have distinct influences on neuroplasticity (Hendrikse et al., 2017). Applying rTMS following exercise may capitalise on the *priming* effects of exercise on neuroplasticity (Hendrikse et al., 2017; Andrews et al., 2020), whereas engaging in exercise following rTMS may have a *consolidating* effect, maintaining an environment conducive to neuroplasticity. For example, performing a bout of exercise performed four hours after learning increases memory retention (van Dongen, Kersten, Wagner, Morris, & Fernández, 2016). Conversely, certain timing/forms of exercise and rTMS may attenuate/block their respective influence on neuroplasticity (see Smith et al., 2018). Our study has provided some indication that higher CRF is associated with greater WM changes following rTMS; however future studies are required to assess the optimal timing of exercise relative to rTMS to maximise effects on neuroplasticity and memory function.

Similar multi-day rTMS protocols to that used in this study have reported increases in functional connectivity (Wang et al., 2014) and associative memory performance persisting for ∼24 hours. In line with these findings, we assessed changes to WM and GABA ∼24 hours after the final rTMS session (see figure 1). This delayed assessment time point (following a night of sleep) may have failed to capture more short-lived effects. Further, to assess changes in GABA concentration, MRS assessments were conducted for the first stimulation condition only. Our assessment of multi-day rTMS on 40 healthy individuals provides one of the larger studies of its type. However, considering the inherent inter-individual variability associated with rTMS (López-Alonso, Cheeran, Río-Rodríguez, & Fernández-Del-Olmo, 2014) our sample size and experimental design (i.e. assessing interactions across timepoints and stimulation and physical activity conditions) may still have lacked experimental power to detect small/moderate effects. Future studies are recommended to explore the interactions between exercise, multi-day rTMS, and GABA concentration with larger sample sizes and within-subject experimental designs.

### Conclusions

In summary, we did not find evidence that multi-day rTMS to parietal cortex influences WM function or GABA concentration. While single doses of rTMS to parietal cortex are associated with transient improvements to aspects of brain structure and function, our results challenge the notion that multiple doses induce longer-lasting effects. However, we do report some evidence of increased WM function and GABA concentration (across voxel sites) following multi-day pre-SMA stimulation. Thus, the pre-SMA may be a useful stimulation target for future studies aimed at enhancing WM function. We also observed positive associations between cardiorespiratory fitness and changes in WM following pre-SMA stimulation, suggesting that habitual exercise may be associated with increased responsiveness to multi-day rTMS. While there are a number of factors known to influence the response to rTMS (Ridding & Ziemann, 2010), increasing cardiorespiratory fitness may provide a novel approach to minimise variability and enhance cognitive outcomes. Given the established clinical utility of both exercise and rTMS, future studies are required to determine whether additive effects may be achieved when applied in tandem.

## Acknowledgements

We also wish to thank the staff at Monash Biomedical Imaging for their assistance with MRI data acquisition.

## Data and material availability

No part of the study procedures or analyses was pre-registered prior to the research being conducted. We report how we determined our sample size, all data exclusions, all inclusion/exclusion criteria, and whether inclusion/exclusion criteria were established prior to data analysis. We also report all manipulations and all measures relevant to our investigation of the effects of multi-day rTMS and cardiorespiratory fitness on working memory and local GABA concentration. However, additional outcome measures were collected in this study, including resting-state fMRI and tests of declarative memory and motor inhibition. These measures were not relevant to the current aims or hypotheses and are therefore not reported. The conditions of our ethical approval do not permit open sharing of participant MRI data without prior informed consent. Therefore, we are unable to publicly archive the raw MRI data used in this study. Readers seeking access to the data should contact the lead author Joshua Hendrikse. Access will be granted to named individuals in accordance with ethical procedures governing the reuse of sensitive data. Specifically, requestors must complete a formal data sharing agreement and make clear the process by which participant consent would be sought. De-identified behavioural data and inputs for group level spectroscopy analyses (i.e. partial volume corrected GABA+ estimates) are available at https://osf.io/y2xm8/, and all code used for data collection and analysis is available at https://github.com/jhendrikse/ex_rtms_code.

## Funding

JH is supported by an Australian government research training scholarship. MY, NR, and JC have all received funding from Monash University, the National Health and Medical Research Council, and the Australian Research Council (ARC). MY has received funding from the Australian Defence Science and Technology (DST), and the Department of Industry, Innovation and Science (DIIS). He has also received philanthropic donations from the David Winston Turner Endowment Fund (which partially supported this study), Wilson Foundation, as well as payment from law firms in relation to court, expert witness, and/or expert review reports. The funding sources had no role in the design, management, data analysis, presentation, or interpretation and write-up of the data.

## Supplementary Materials

### Comparison of working memory performance following the first week of stimulation

To account for the possibility of carry-over effects between stimulation conditions, we assessed working memory following the first stimulation condition only (i.e. between-subject factors of stimulation site, and physical activity group).

For reaction time, there was no significant main effect of Time (baseline, post-rTMS; *F*_1,34_ = 0.44, p = .51, *η*^2^*_p_* = .01) or Physical Activity (high PA, low PA; *F*_1,34_ = 0.62, p = .44, *η*^2^*_p_*= .02). However, there was a significant main effect of Stimulation Condition (parietal, pre-SMA; *F*_1,34_ = 5.51, p = .025, *η*^2^*_p_*= .14, BF_10_ = 2.55), due to faster reaction time in the pre-SMA stimulation condition (averaged over Time and Physical Activity) (M = 0.53, SE = 0.02), relative to the parietal stimulation condition (M = 0.59, SE = 0.02), with a small effect size (Cohen’s d = 0.38; p = .025 [Tukey corrected post hoc test], BF_10_ = 2.55). There was also a significant interaction between Stimulation Condition and Time (*F*_1,34_ = 7.36, p = .01, *η*^2^*_p_* = .18, BF_10_ = 5.32), due to a faster reaction time post pre-SMA stimulation (post pre-SMA, M = 0.52, SE = 0.02), relative to post parietal stimulation (post parietal, M = 0.60, SE = 0.02), with a small effect size (Cohen’s d = 0.49; p = .025 [Tukey corrected post hoc test], BF_10_ = 5.32). However, no significant interaction was observed between Stimulation Condition, Time, and Physical Activity (*F*_1,34_ = 0.92, p = .34, *η*^2^*_p_* = .03).

For *d’*, there was no significant main effect of Stimulation Condition (parietal, pre-SMA; *F*_1,34_ = 0.72, p = .40, *η*^2^*_p_* = .02), or Physical Activity (high PA, low PA; *F*_1,34_ = 0.08, p = .78, *η*^2^*_p_* = .00). However, there was a significant main effect of Time (baseline, post-rTMS; *F*_1,34_ = 4.83, p = .03, *η*^2^*_p_* = .12), due to higher *d’* values (averaged over Stimulation Condition and Physical Activity) post rTMS (M = 3.10, SE = 0.13), relative to baseline (M = 2.92, SE = 0.13), with a small effect size (Cohen’s d = 0.33; p = .035 [Tukey corrected post hoc test], BF_10_ = 1.37). There were also significant interactions between Stimulation Condition and Time (*F*_1,34_ = 4.67, p = .04, *η*^2^*_p_* = .12, BF_10_ = 1.60), and Time and Physical Activity (*F*_1,34_ = 4.26, p = .047, *η*^2^*_p_* = .11, BF_10_ = 1.41). Tukey corrected post-hoc tests revealed a significant increase in *d’* values across time in the high PA group with a small effect size (baseline M = 2.88, SE = 0.18; post-rTMS M = 3.22, SE = 0.18, p = .024, Cohen’s d = 0.49, BF_10_ = 1.41), and a significant increase following pre-SMA stimulation, with a medium effect size (baseline M = 2.73, SE = 0.16; post-rTMS M = 3.08, SE = 0.17; p = .020, Cohen’s d = 0.50, BF_10_ = 1.60). However, no significant interaction was observed between Stimulation Condition, Time, and Physical Activity (*F*_1,34_ = 0.02, p = .89, *η*^2^*_p_* = .00).

### Follow-up assessment of working memory performance

To assess whether stimulation induced lasting effects on working memory performance, working memory assessments taken at follow up (∼1.5 weeks following final rTMS session) were compared between stimulation groups using an independent ANOVA. No significant differences in reaction time (s) were found between parietal stimulation and pre-SMA stimulation (*F*_1,34_ = 3.15, p = .085, *η*^2^*_p_* = .08), nor between high PA and low PA (*F*_1,34_ = 0.08, p = .79, *η*^2^*_p_* = .00). There was no significant interaction between Stimulation Condition and Activity Group (*F*_1,34_ = 0.11, p = .74, *η*^2^*_p_* = .00). Similarly, no significant differences in *d’* values were found between parietal stimulation and pre-SMA stimulation (*F*_1,34_ = 1.20, p = .28, *η*^2^*_p_* = .03), nor between high PA and low PA (*F*_1,34_ = 1.56, p = .22, *η*^2^*_p_* = .04). There was no significant interaction between Stimulation Condition and Activity Group (*F*_1,34_ = 1.75, p = .19, *η*^2^*_p_* = .05). Overall, these results suggest that multi-day rTMS did not exert a long-lasting influence on working memory performance.

### MRS voxel localisation

Across stimulation conditions and physical activity groups, the overlap in voxel position (mean ± SD) for baseline and post-rTMS assessments for the parietal voxel was 88.2% ± 5.7% and 90.1% ± 5.2% for the pre-SMA voxel. A repeated-measures ANOVA conducted on proportion of baseline and post-rTMS voxel overlap showed no significant showed no significant main effect of Voxel Site (parietal, pre-SMA; *F*_1,33_ = 2.80, p = .10, *η*^2^*_p_* = .08), Stimulation Condition (parietal, pre-SMA; *F*_1,33_ = 0.20, p = .66, *η*^2^*_p_* = .00), or Physical Activity (high PA, low PA; *F*_1,33_ = 0.72, p = .40, *η*^2^*_p_* = .02), nor interaction between Voxel Site and Stimulation Condition (*F*_1,33_ = 0.13, p = .72, *η*^2^*_p_* = .00), or Voxel Site, Stimulation Condition, and Physical Activity (*F*_1,33_ = 0.40, p = .53, *η*^2^*_p_* = .01).

### Analysis of MRS voxel grey matter, white matter, and CSF fractions

We assessed the proportion of grey matter, white matter, and CSF across voxels, time, and stimulation conditions.

### Grey matter

A repeated-measures ANOVA was conducted on the proportion of GM within voxels, and showed no main effect of Time (baseline, post-rTMS; *F*_1,34_ = 0.14, p = .71, *η*^2^*_p_* = .00), Stimulation Condition (parietal, pre-SMA; *F*_1,34_ = 0.65, p = .42, *η*^2^*_p_* = .02), Physical Activity (high PA, low PA; *F*_1,34_ = 3.60, p = .066, *η*^2^*_p_* = .10), or Voxel Site (parietal, pre-SMA; *F*_1,34_ = 2.42, p = .13, *η*^2^*_p_* = .07), nor interaction between Stimulation Condition and Time (*F*_1,34_ = 0.01, p = .94, *η*^2^*_p_* = .00), Stimulation Condition, Time, and Physical Activity (*F*_1,34_ = 0.68, p = .42, *η*^2^*_p_* = .02) or Stimulation Condition, Time, Physical Activity, and Voxel Site (*F*_1,34_ = 0.17, p = .68, *η*^2^*_p_* = .01).

### White matter

A repeated-measures ANOVA was conducted on the proportion of WM within voxels, and showed no main effect of Time (baseline, post-rTMS; *F*_1,34_ = 0.14, p = .71, *η*^2^*_p_* = .00) or Stimulation Condition (parietal, pre-SMA; *F*_1,34_ = 0.51, p = .48, *η*^2^*_p_* = .01). However, there was a significant main effect of Physical Activity (high PA, low PA; *F*_1,34_ = 4.32, p = .045, *η*^2^*_p_* = .11) with significantly higher WM proportions in high PA subjects (M = 46.2%, SE = 1.3%), relative to low PA subjects (M = 42.3%, SE = 1.3%). Additionally, we observed a main effect of Voxel Site (*F*_1,34_ = 16.72, p = .0003, *η*^2^*_p_* = .33), with significantly higher WM proportions in the parietal (M = 46.8%, SE = 1.1%) voxel relative to pre-SMA (M = 41.7%, SE = 1.1%). There was no significant interaction between Stimulation Condition and Time (*F*_1,34_ = 0.01, p = .92, *η*^2^*_p_* = .00), Stimulation Condition, Time, and Physical Activity (*F*_1,34_ = 0.74, p = .40, *η*^2^*_p_* = .02), or Stimulation Condition, Time, Physical Activity, and Voxel Site (*F*_1,34_ = 0.35, p = .56, *η*^2^*_p_* = .01).

### CSF

A repeated-measures ANOVA was conducted on the proportion of WM within voxels, and showed no main effect of Time (baseline, post-rTMS; *F*_1,34_ = 0.04, p = .84, *η*^2^*_p_* = .00), Stimulation Condition (parietal, pre-SMA; *F*_1,34_ = 0.03, p = .87, *η*^2^*_p_* = .00), or Physical Activity (high PA, low PA; *F*_1,34_ = 0.68, p = .41, *η*^2^*_p_* = .02). However, we observed a main effect of Voxel Site (*F*_1,34_ = 14.71, p = .0005, *η*^2^*_p_* = .30), with significantly higher CSF proportions in the pre-SMA voxel (M = 15.9%, SE = 0.8%) voxel relative to parietal voxel (M = 12.3%, SE = 0.8%). There was no significant interaction between Stimulation Condition and Time (*F*_1,34_ = 0.03, p = .87, *η*^2^*_p_* = .00), Stimulation Condition, Time, and Physical Activity (*F*_1,34_ = 0.60, p = .45, *η*^2^*_p_* = .02), or Stimulation Condition, Time, Physical Activity, and Voxel Site (*F*_1,34_ = 0.28, *p* = .60, *η*^2^*_p_* = .01).

In summary, there were also no significant differences in the proportion of grey matter, white matter, and CSF fractions within voxels across time and stimulation conditions. Between parietal and pre-SMA voxels there was a significant difference in white matter (Parietal M = 46.8 %, SE = 1.1%; pre-SMA M = 41.7 %, SE = 1.1 %) and CSF (Parietal M = 12.3 %, SE = 0.8%; pre-SMA M = 15.9 %, SE = 0.8 %). There was also a significant main effect of Physical Activity on white matter proportion.

### Assessment of MRS data quality

To examine the consistency of GABA+ quantification across time, stimulation conditions, voxel sites and physical activity groups, a repeated-measures ANOVA was conducted on E_GABA_ and GABA FWHM values.

For E_GABA_, there was no main effect of Time (baseline, post-rTMS; *F*_1,33_ = 0.28, p = .60, *η*^2^*_p_* = .01), Stimulation Condition (parietal, pre-SMA; *F*_1,33_ = 0.81, p = .37, *η*^2^*_p_* = .02), or Physical Activity (high PA, low PA; *F*_1,33_ = 3.39, p = .074, *η*^2^*_p_* = .09), nor interaction between Stimulation Condition and Time (*F*_1,33_ = 0.10, p = .76, *η*^2^*_p_* = .00), or Stimulation Condition, Time, and Physical Activity (*F*_1,33_ = 0.51, p = .48, *η*^2^*_p_* = .02). However, there was a significant main effect of Voxel Site (*F*_1,33_ = 21.52, p = .0001, *η*^2^*_p_* = .39), with higher average E_GABA_ values in the pre-SMA voxel (M = 11.59, SE = 0.33) relative to parietal voxel site (M = 9.54, SE = 0.33). A post-hoc with Bonferroni correction determined that this difference was significant (p = 0.00005, Cohen’s d = 0.76, BF_10_ = 16286.84). There was no significant interaction between Voxel Site, Stimulation Condition, Time, and Physical Activity (*F*_1,33_ = 0.21, p = .65, *η*^2^*_p_* = .01).

Similarly, analysis of GABA FWHM values revealed no main effect of Time (baseline, post-rTMS; *F*_1,34_ = 0.02, p = .90, *η*^2^*_p_* = .00), Stimulation Condition (parietal, pre-SMA; *F*_1,34_ = 0.16,p = .69, *η*^2^*_p_* = .00), or Physical Activity (high PA, low PA; *F*_1,34_ = 1.84, p = .18, *η*^2^*_p_* = .05), nor interaction between Stimulation Condition and Time (*F*_1,34_ = 0.72, p = .40, *η*^2^*_p_* = .02), or Stimulation Condition, Time, and Physical Activity (*F*_1,34_ = 0.20, p = .66, *η*^2^*_p_* = .01). However, there was a significant main effect of Voxel Site (*F*_1,34_ = 25.34, p = .00002, *η*^2^*_p_* = .43) with higher average FWHM values in the pre-SMA voxel (M = 24.81, SE = 0.95), relative to parietal voxel (M = 18.28, SE = 0.95). A post-hoc test with Bonferroni correction determined that this difference was significant (p = .00002, Cohen’s d = .82, BF_10_ = 169200.10). There was no significant interaction between Voxel Site, Stimulation Condition, Time, and Physical Activity (*F*_1,34_ = 0.48, p = .49, *η*^2^*_p_* = .01).

GABA+ estimates were reported relative to the unsuppressed water signal. Therefore, we also assessed the stability of water estimates using FWHM values, across baseline and post-rTMS assessments, voxel positions (parietal, pre-SMA), and between stimulation and physical activity groups. There was no significant main effect of Time (baseline, post-rTMS; *F*_1,34_ = 0.11, p = .74, *η*^2^*_p_* = .00), Stimulation Condition (parietal, pre-SMA; *F*_1,34_ = 0.05, p = .82, *η*^2^*_p_* = .00), Physical Activity (high PA, low PA; *F*_1,34_ = 0.02, p = .89, *η*^2^*_p_* = .00), or Voxel Site (parietal, pre-SMA; *F*_1,34_ = 3.56, p = .068, *η*^2^*_p_* = .09). There was also no significant interaction between Stimulation Condition and Time (*F*_1,34_ = 1.05, p = .31, *η*^2^*_p_* = .03), Stimulation Condition, Time, and Physical Activity (*F*_1,34_ = 0.01, p = .92, *η*^2^*_p_* = .00), or Voxel Site, Stimulation Condition, Time, Physical Activity (*F*_1,34_ = 0.01, p = .93, *η*^2^*_p_* = .00).

Overall, our results demonstrate consistent data quality within each voxel location across baseline and post-rTMS assessments, stimulation conditions, and physical activity groups. We did observe evidence of differences in data quality between voxel locations, with higher E_GABA_ and GABA FWHM in the pre-SMA voxel, relative to the parietal voxel location. These differences may be partially attributable to differences in WM and CSF tissue fractions between parietal and pre-SMA voxels.

### Analysis of baseline GABA+ concentration between parietal and pre-SMA voxels

To account for the possibility that regional differences in baseline GABA+ concentration may have influenced the response to rTMS, we conducted Wilcoxon signed-ranks test on baseline GABA+ concentrations between parietal and pre-SMA voxels across participants. There were no significant differences in GABA+ concentration between parietal (M = 2.50, SD = 0.53) and pre-SMA (M = 2.83, SD = 1.42) voxel sites (W = 305, p = .67, r = .08, BF_10_ = 0.45), suggesting that regional differences in GABA concentration are unlikely to have influenced the response to stimulation.

### Analysis of individual change in GABA+ concentration following stimulation

To account for variability in baseline estimates between stimulation conditions, we also conducted a 2 x 2 x 2 repeated-measures ANOVA on each subject’s change in GABA+ concentration at each voxel location (post-rTMS GABA – baseline GABA). There was no main effect of Voxel Site (parietal, pre-SMA; *F*_1,31_ = 0.33, p = .57, *η*^2^*_p_* = .01), or Physical Activity (high PA, low PA; *F*_1,31_ = 0.00, p = .97, *η*^2^*_p_* = .00). A significant main effect of Stimulation Condition was observed (parietal, pre-SMA; *F*_1,31_ = 5.26, p = .029, *η*^2^*_p_* = .15), with a significant difference in Δ GABA+ estimates (averaged across voxel locations) between stimulation conditions with a medium effect size (pre-SMA M = 0.60, SE = 0.22; parietal M = -0.11, SE = 0.22; Cohen’s d = .39; p = .029 [Tukey corrected post hoc test], BF_10_ = 1.07. We also observed a significant interaction between Stimulation Condition and Physical Activity (*F*_1,31_ = 4.33, p = .046, *η*^2^*_p_* = .12), suggesting that the significant difference in Δ GABA+ estimates between stimulation conditions occurred in the low PA group, with a medium effect size (pre-SMA M = 0.92, SE = 0.31; parietal M = -0.43, SE = 0.32; Cohen’s d = 0.52; p = .028 [Tukey corrected post hoc test], BF_10_ = 1.05. There was no significant interaction between Stimulation Condition and Voxel Site (*F*_1,31_ = 0.95, p = .34, *η*^2^*_p_* = .03). However, we did observe a significant interaction between Voxel Site, Stimulation Condition, and Physical Activity (*F*_1,31_ = 4.65, p = .039, *η*^2^*_p_* = .13, BF_10_ = 4.01). A post-hoc Tukey’s test revealed a significant difference in Δ pre-SMA GABA+ concentration between stimulation conditions, in low PA individuals with a medium effect size (parietal rTMS M = -0.97, SE = 1.60; pre-SMA rTMS M = 1.45, SE = 2.21; p = .011, Cohen’s d = 0.62, BF_10_ = 4.01), with no other significant differences observed (all Cohen’s d < .44, p > .18).

**Figure S1.**
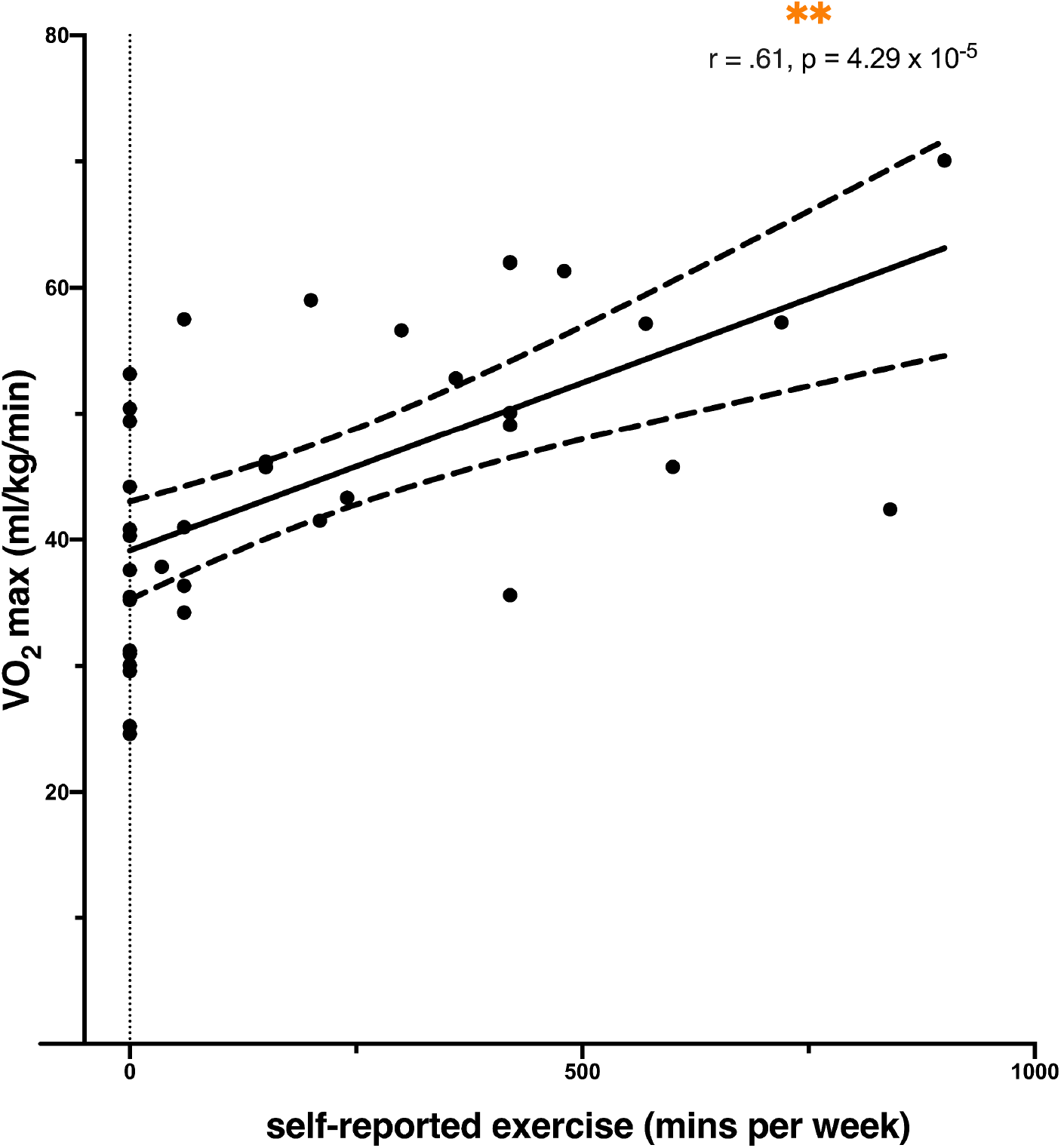
Association between self-reported exercise and VO_2_max. There was a significant positive association observed between self-reported exercise and VO_2_max (across PA groups, r = .61, p = 4.29 x 10^-5^). Circles represent individual scores, solid line shows regression line of best fit, dotted lines depict 95% confidence interval. VO_2_max score is represented on the y-axis and exercise level (self-reported weekly minutes of exercise) is represented on the x-axis. ** p < 5 × 10^-5^.

**Figure S2.**
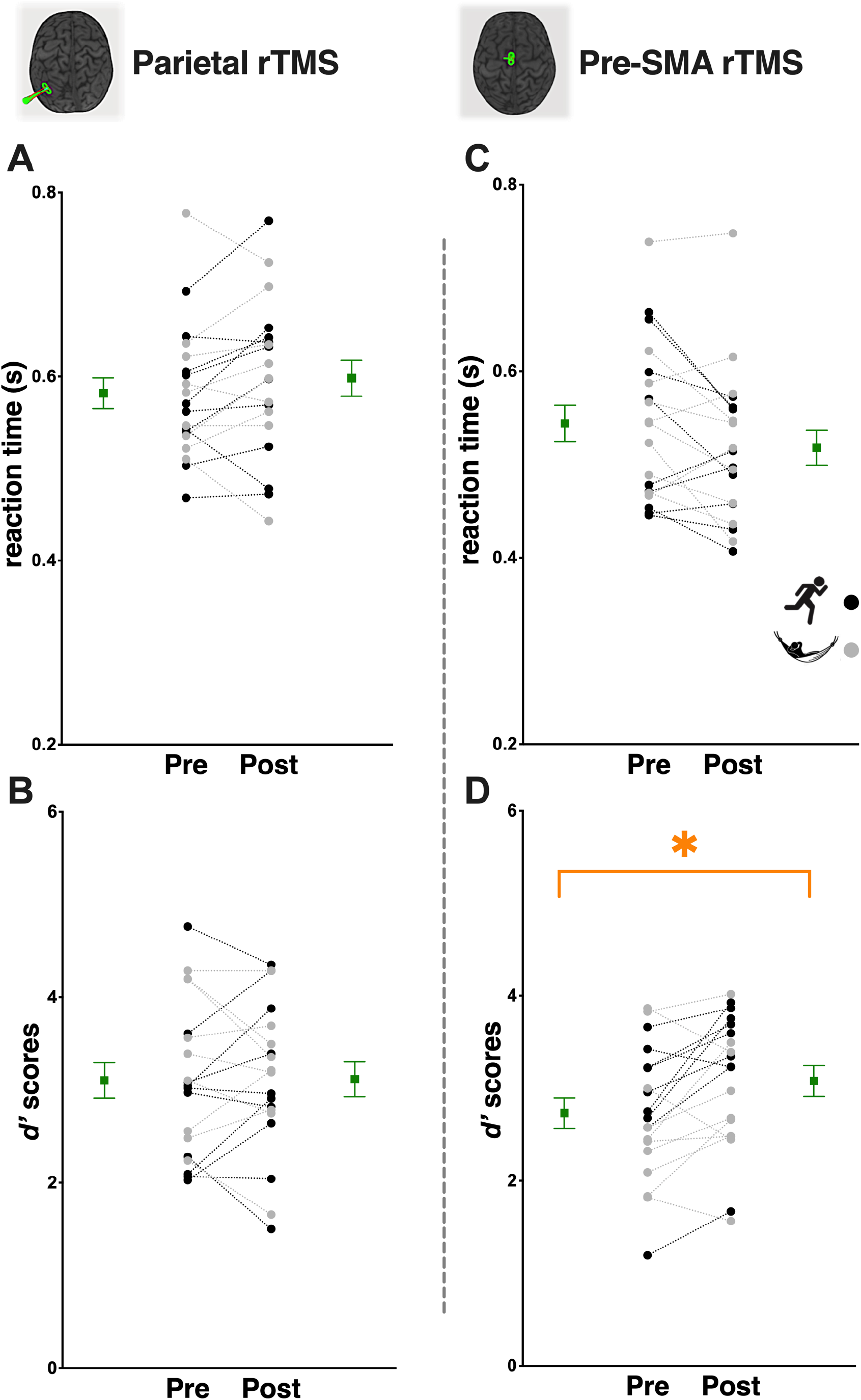
Effects of multi-day rTMS and physical activity on working memory following the first stimulation condition only. Some evidence of improved working memory performance following multi-day pre-SMA rTMS; no change following multi-day parietal rTMS. The left panel **(A,B)** shows performance on the n-back task following parietal rTMS (**A,** reaction time; **B,** *d’* scores). The right panel **(C,D)** shows performance on the n-back task following pre-SMA rTMS (**C,** reaction time; **D,** *d’* scores). Circles represent individual scores with high PA individuals depicted in black and low PA individuals in grey; squares and error bars depict group mean and standard error. * p < .05.

**Figure S3.**
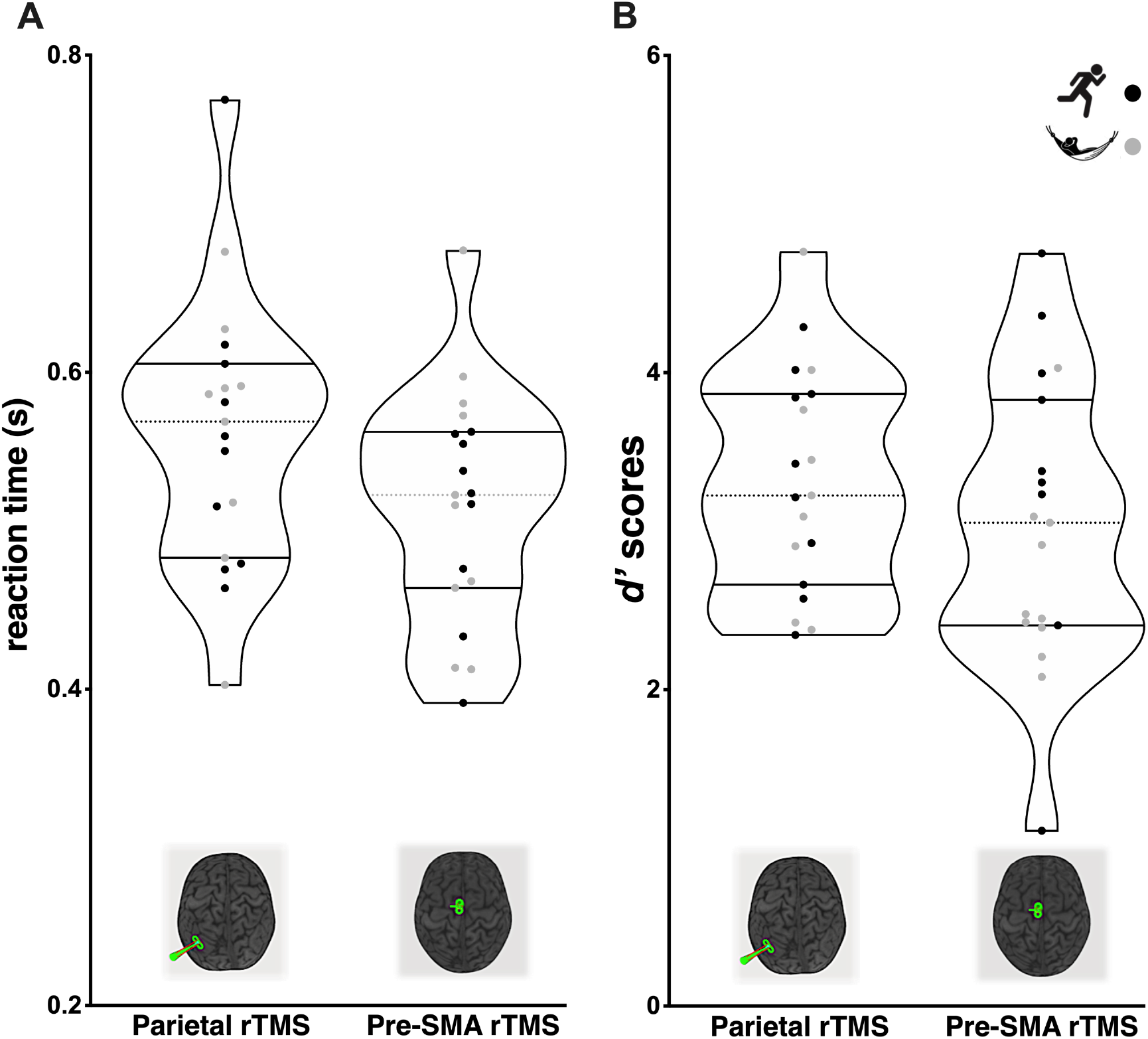
Follow-up assessment of working memory function. No evidence of long-lasting changes (∼15 days) to working memory performance following multi-day rTMS. **(A)** Reaction time at follow-up assessment (parietal stimulation, left; pre-SMA stimulation, right); **B)** *d’* values at follow-up assessment (parietal stimulation, left; pre-SMA stimulation, right). Circles represent individual scores, with high PA individuals depicted in black and low PA individuals in grey. For each group, mean values are indicated by horizontal dashed lines, whereas higher and lower quartiles are indicated by dotted lines.

**Figure S4.**
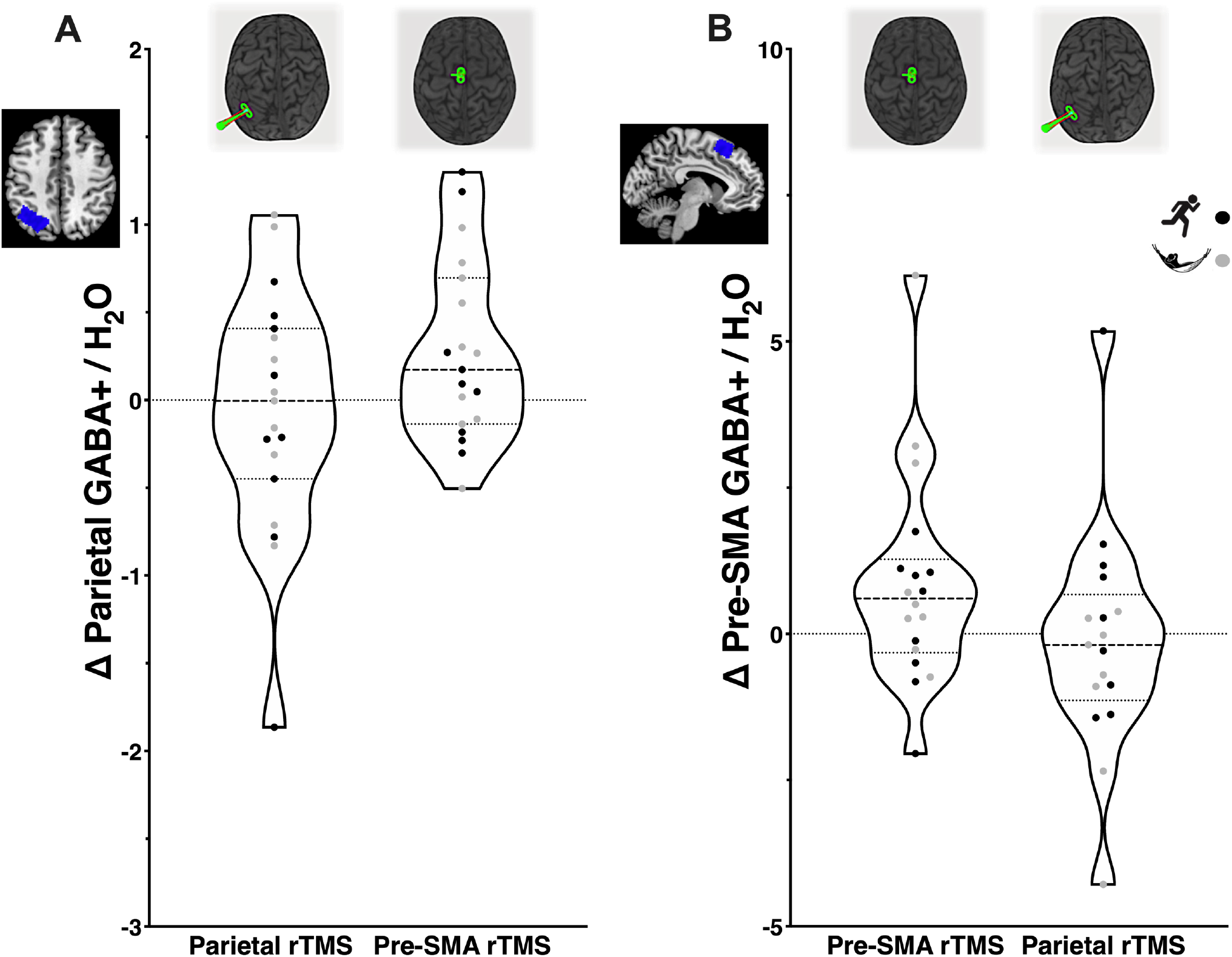
Individual change in GABA+ concentration following multi-day rTMS. Evidence of a generalised increase in GABA+ concentration following multi-day pre-SMA rTMS. **(A)** GABA+ estimates (post-pre) in the parietal voxel following multi-day rTMS (parietal stimulation, left; SMA stimulation, right). **B)** GABA+ estimates (post-pre) in the pre-SMA voxel following multi-day rTMS (pre-SMA stimulation, left; parietal stimulation, right). Circles represent individual scores, with high PA individuals depicted in black and low PA individuals in grey. For each group, mean values are indicated by horizontal dashed lines, whereas higher and lower quartiles are indicated by dotted lines.

**Table S1:** Comparison of demographic variables and baseline WM performance between week one stimulation groups

| Demographic variable | Mean (SD)<br>Parietal rTMS | Mean (SD)<br>Pre-SMA rTMS | Statistic | df | p | Effect size |
| --- | --- | --- | --- | --- | --- | --- |
| Age (years) | 26.26 (10.64) | 24.84 (8.65) | 0.45<sup>a</sup> | 1,36 | .65 | 0.15<sup>b</sup> |
| Sex | 47.4 % female | 63.2 % female | 0.96<sup>c</sup> | 1 | .33 | .16<sup>d</sup> |
| Baseline reaction time | 0.58 (0.07) | 0.54 (0.08) | 1.48<sup>a</sup> | 1,36 | .15 | 0.48<sup>b</sup> |
| Baseline $d'$ | 3.10 (0.83) | 2.73 (0.72) | 1.48<sup>a</sup> | 1,36 | .15 | 0.48<sup>b</sup> |

<sup>a</sup> $t$ statistic (Independent-samples $t$ -test); <sup>b</sup> Cohen's $d$ ; <sup>c</sup> chi-square statistic; <sup>d</sup> Cramer's $V$

## Notes

### Competing Interest Statement

The authors have declared no competing interest.

## References

1. Albouy, P., Weiss, A., Baillet, S., & Zatorre, R. J. (2017). Selective Entrainment of Theta Oscillations in the Dorsal Stream Causally Enhances Auditory Working Memory Performance. Neuron, 94(1), 193–206.e5. https://doi.org/10.1016/j.neuron.2017.03.015

2. Baddeley, A. (1992). Working memory. Science, 255(5044), 556–559. https://doi.org/10.1126/science.1736359

3. Baddeley, A. (2010). Working memory. Current Biology, 20(4), 136–140. https://doi.org/10.1016/j.cub.2009.12.014

4. Baddeley, A. D., Bressi, S., Della Sala, S., Logie, R., & Spinnler, H. (1991). The decline of working memory in alzheimer’s disease: A longitudinal study. Brain, 114(6), 2521–2542. https://doi.org/10.1093/brain/114.6.2521

5. Bagherzadeh, Y., Khorrami, A., Zarrindast, M. R., Shariat, S. V., & Pantazis, D. (2016). Repetitive transcranial magnetic stimulation of the dorsolateral prefrontal cortex enhances working memory. Experimental Brain Research. https://doi.org/10.1007/s00221-016-4580-1

6. Barr, M. S., Farzan, F., Arenovich, T., Chen, R., Fitzgerald, P. B., & Daskalakis, Z. J. (2011). The effect of repetitive transcranial magnetic stimulation on gamma oscillatory activity in schizophrenia. PLoS ONE, 6(7). https://doi.org/10.1371/journal.pone.0022627

7. Borg, G. A. V. (1982). Psychophysical bases of perceived exertion. Medicine and Science in Sports and Exercise.

8. Brunoni, A. R., & Vanderhasselt, M. A. (2014). Working memory improvement with non-invasive brain stimulation of the dorsolateral prefrontal cortex: A systematic review and meta-analysis. Brain and Cognition, 86(1), 1–9. https://doi.org/10.1016/j.bandc.2014.01.008

9. Buchsbaum, B. R., & D’Esposito, M. (2019, March 1). A sensorimotor view of verbal working memory. Cortex. Masson SpA. https://doi.org/10.1016/j.cortex.2018.11.010

10. Castro-Alamancos, M. A., Donoghue, J. P., & Connors, B. W. (1995). Different forms of synaptic plasticity in somatosensory and motor areas of the neocortex. Journal of Neuroscience, *15*(7 II), 5324–5333. https://doi.org/10.1523/jneurosci.15-07-05324.1995

11. Casula, E. P., Pellicciari, M. C., Picazio, S., Caltagirone, C., & Koch, G. (2016). Spike-timing-dependent plasticity in the human dorso-lateral prefrontal cortex. NeuroImage, 143, 204–213. https://doi.org/10.1016/j.neuroimage.2016.08.060

12. Chung, S. W., Lewis, B. P., Rogasch, N. C., Saeki, T., Thomson, R. H., Hoy, K. E., … Fitzgerald, P. B. (2017). Demonstration of short-term plasticity in the dorsolateral prefrontal cortex with theta burst stimulation: A TMS-EEG study. Clinical Neurophysiology, 128(7), 1117–1126. https://doi.org/10.1016/j.clinph.2017.04.005

13. Cirillo, J., Lavender, A. P., Ridding, M. C., & Semmler, J. G. (2009). Motor cortex plasticity induced by paired associative stimulation is enhanced in physically active individuals. The Journal of Physiology, 587(Pt 24), 5831–5842. https://doi.org/10.1113/jphysiol.2009.181834

14. Coxon, J. P., Cash, R. F. H., Hendrikse, J. J., Rogasch, N. C., Stavrinos, E., Suo, C., & Yücel, M. (2018). GABA concentration in sensorimotor cortex following high-intensity exercise and relationship to lactate levels. Journal of Physiology, 596(4), 691–702. https://doi.org/10.1113/JP274660

15. Craig, C. L., Marshall, A. L., Sjöström, M., Bauman, A. E., Booth, M. L., Ainsworth, B. E., … Oja, P. (2003). International Physical Activity Questionnaire: 12-Country Reliability and Validity. Medicine & Science in Sports & Exercise, 35(8), 1381–1395. https://doi.org/10.1249/01.MSS.0000078924.61453.FB

16. Daskalakis, Z. J., Moller, B., Christensen, B. K., Fitzgerald, P. B., Gunraj, C., & Chen, R. (2006). The effects of repetitive transcranial magnetic stimulation on cortical inhibition in healthy human subjects. Exp Brain Res, 174(3), 403–412. https://doi.org/10.1007/s00221-006-0472-0

17. Di Martino, A., Scheres, A., Margulies, D. S., Kelly, A. M. C., Uddin, L. Q., Shehzad, Z., … Milham, M. P. (2008). Functional connectivity of human striatum: A resting state fMRI study. Cerebral Cortex, 18(12), 2735–2747. https://doi.org/10.1093/cercor/bhn041

18. Edden, R. A. E., Puts, N. A. J., Harris, A. D., Barker, P. B., & C. John Evans. (2014). Gannet: A Batch-Processing Tool for the Quantitative Analysis of Gamma-Aminobutyric Acid–Edited MR Spectroscopy Spectra Richard, 40(6), 1445–1452.

19. Edden, R. A. E., Puts, N. A. J., Harris, A. D., Barker, P. B., & Evans, C. J. (2014). Gannet: A batch-processing tool for the quantitative analysis of gamma-aminobutyric acid-edited MR spectroscopy spectra. Journal of Magnetic Resonance Imaging, 40(6), 1445–1452. https://doi.org/10.1002/jmri.24478

20. Evans, C. J., McGonigle, D. J., & Edden, R. A. E. (2010). Diurnal stability of γ-aminobutyric acid concentration in visual and sensorimotor cortex. Journal of Magnetic Resonance Imaging, 31(1), 204–209. https://doi.org/10.1002/jmri.21996

21. Fitzgerald, P. B., Fountain, S., & Daskalakis, Z. J. (2006). A comprehensive review of the effects of rTMS on motor cortical excitability and inhibition. Clinical Neurophysiology, 117(12), 2584–2596. https://doi.org/10.1016/j.clinph.2006.06.712

22. Gaudeau-Bosma, C., Moulier, V., Allard, A. C., Sidhoumi, D., Bouaziz, N., Braha, S., … Januel, D. (2013). Effect of two weeks of rTMS on brain activity in healthy subjects during an n-back task: A randomized double blind study. Brain Stimulation, 6(4), 569–575. https://doi.org/10.1016/j.brs.2012.10.009

23. Greenhouse, I., Noah, S., Maddock, R. J., & Ivry, R. B. (2016). Individual differences in GABA content are reliable but are not uniform across the human cortex. NeuroImage, 139, 1–7. https://doi.org/10.1016/j.neuroimage.2016.06.007

24. Guiney, H., & Machado, L. (2013, December 11). Benefits of regular aerobic exercise for executive functioning in healthy populations. Psychonomic Bulletin and Review. Springer. https://doi.org/10.3758/s13423-012-0345-4

25. Guse, B., Falkai, P., Gruber, O., Whalley, H., Gibson, L., Hasan, A., … Wobrock, T. (2013). The effect of long-term high frequency repetitive transcranial magnetic stimulation on working memory in schizophrenia and healthy controls-A randomized placebo-controlled, double-blind fMRI study. Behavioural Brain Research, 237(1), 300–307. https://doi.org/10.1016/j.bbr.2012.09.034

26. Hallett, M. (2007). Transcranial Magnetic Stimulation: A Primer. Neuron, 55(2), 187–199. https://doi.org/10.1016/j.neuron.2007.06.026

27. Hamada, M., Ugawa, Y., & Tsuji, S. (2009). High-frequency rTMS over the supplementary motor area improves bradykinesia in Parkinson’s disease: subanalysis of double-blind sham-controlled study. Journal of the Neurological Sciences, 287(1–2), 143–146. https://doi.org/10.1016/j.jns.2009.08.007

28. Hamidi, M., Tononi, G., & Postle, B. R. (2008). Evaluating frontal and parietal contributions to spatial working memory with repetitive transcranial magnetic stimulation. Brain Research, 1230, 202–210. https://doi.org/10.1016/j.brainres.2008.07.008

29. Hendrikse, J., Coxon, J., Thompson, S., Suo, C., Fornito, A., Yücel, M., & Rogasch, N. C. (2020). Multi-day rTMS exerts site-specific effects on functional connectivity but does not influence associative memory performance. BioRxiv, 2020.04.23.056655. https://doi.org/10.1101/2020.04.23.056655

30. Hendrikse, J., Kandola, A., Coxon, J., Rogasch, N., & Yücel, M. (2017). Combining aerobic exercise and repetitive transcranial magnetic stimulation to improve brain function in health and disease. Neuroscience & Biobehavioral Reviews, 83(September), 11–20. https://doi.org/10.1016/j.neubiorev.2017.09.023

31. Henry, M. E., Lauriat, T. L., Shanahan, M., Renshaw, P. F., & Jensen, J. E. (2011). Accuracy and stability of measuring GABA, glutamate, and glutamine by proton magnetic resonance spectroscopy: A phantom study at 4 Tesla. Journal of Magnetic Resonance, 208(2), 210–218. https://doi.org/10.1016/j.jmr.2010.11.003

32. Hikosaka, O., Sakai, K., Miyauchi, S., Takino, R., Sasaki, Y., & Pütz, B. (1996). Activation of human presupplementary motor area in learning of sequential procedures: A functional MRI study. Journal of Neurophysiology, 76(1), 617–621. https://doi.org/10.1152/jn.1996.76.1.617

33. Hikosaka, Okihide, Nakamura, K., Sakai, K., & Nakahara, H. (2002). Central mechanisms of motor skill learning. Current Opinion in Neurobiology, 12(2), 217–222. https://doi.org/https://doi.org/10.1016/S0959-4388(02)00307-0

34. Hoogendam, J. M., Ramakers, G. M. J., & Di Lazzaro, V. (2010). Physiology of repetitive transcranial magnetic stimulation of the human brain. Brain Stimulation, 3(2), 95–118. https://doi.org/10.1016/j.brs.2009.10.005

35. Hoy, K. E., Bailey, N., Michael, M., Fitzgibbon, B., Rogasch, N. C., Saeki, T., & Fitzgerald, P. B. (2016). Enhancement of working memory and task-related oscillatory activity following intermittent theta burst stimulation in healthy controls. Cerebral Cortex, 26(12), 4563–4573. https://doi.org/10.1093/cercor/bhv193

36. Irlbacher, K., Kraft, A., Kehrer, S., & Brandt, S. A. (2014, October 1). Mechanisms and neuronal networks involved in reactive and proactive cognitive control of interference in working memory. Neuroscience and Biobehavioral Reviews. Elsevier Ltd. https://doi.org/10.1016/j.neubiorev.2014.06.014

37. Ishihara, T., Miyazaki, A., Tanaka, H., & Matsuda, T. (2020). Identification of the brain networks that contribute to the interaction between physical function and working memory: an fMRI investigation with over 1,000 healthy adults. NeuroImage, 117152. https://doi.org/10.1016/j.neuroimage.2020.117152

38. Jenkinson, M., Beckmann, C. F., Behrens, T. E. J., Woolrich, M. W., & Smith, S. M. (2012). FSL. NeuroImage, 62, 982–790. https://doi.org/10.1016/j.neuroimage.2011.09.015

39. Kandola, A., Hendrikse, J., Lucassen, P. J., & Yücel, M. (2016). Aerobic exercise as a tool to improve hippocampal plasticity and function in humans: Practical implications for mental health treatment. Frontiers in Human Neuroscience, 10(JULY2016). https://doi.org/10.3389/fncel.2016.00373

40. Lett, T. A., Voineskos, A. N., Kennedy, J. L., Levine, B., & Daskalakis, Z. J. (2014). Treating working memory deficits in schizophrenia: A review of the neurobiology. Biological Psychiatry, 75(5), 361–370. https://doi.org/10.1016/j.biopsych.2013.07.026

41. López-Alonso, V., Cheeran, B., Río-Rodríguez, D., & Fernández-Del-Olmo, M. (2014a). Inter-individual variability in response to non-invasive brain stimulation paradigms. Brain Stimulation, 7(3), 372–380. https://doi.org/10.1016/j.brs.2014.02.004

42. López-Alonso, V., Cheeran, B., Río-Rodríguez, D., & Fernández-Del-Olmo, M. (2014b). Inter-individual variability in response to non-invasive brain stimulation paradigms. Brain Stimulation, 7(3), 372–380. https://doi.org/10.1016/j.brs.2014.02.004

43. Luber, B., Kinnunen, L. H., Rakitin, B. C., Ellsasser, R., Stern, Y., & Lisanby, S. H. (2007). Facilitation of performance in a working memory task with rTMS stimulation of the precuneus: Frequency- and time-dependent effects. Brain Research, 1128(1), 120–129. https://doi.org/10.1016/j.brainres.2006.10.011

44. Ly, A., Verhagen, J., & Wagenmakers, E. J. (2016). Harold Jeffreys’s default Bayes factor hypothesis tests: Explanation, extension, and application in psychology. Journal of Mathematical Psychology, 72, 19–32. https://doi.org/10.1016/j.jmp.2015.06.004

45. Maddock, R. J., Casazza, G. A., Fernandez, D. H., & Maddock, M. I. (2016). Acute Modulation of Cortical Glutamate and GABA Content by Physical Activity. Journal of Neuroscience, 36(8), 2449–2457. https://doi.org/10.1523/JNEUROSCI.3455-15.2016

46. Maddock, Richard J., & Buonocore, M. H. (2012). MR Spectroscopic studies of the brain in psychiatric disorders. Current Topics in Behavioral Neurosciences, 11, 199–251. https://doi.org/10.1007/7854_2011_197

47. Marvel, C. L., Morgan, O. P., & Kronemer, S. I. (2019, July 1). How the motor system integrates with working memory. Neuroscience and Biobehavioral Reviews. Elsevier Ltd. https://doi.org/10.1016/j.neubiorev.2019.04.017

48. Matsunaga, K., Maruyama, A., Fujiwara, T., Nakanishi, R., Tsuji, S., & Rothwell, J. C. (2005). Increased corticospinal excitability after 5 Hz rTMS over the human supplementary motor area. Journal of Physiology, 562(1), 295–306. https://doi.org/10.1113/jphysiol.2004.070755

49. McDonnell, M. N., Buckley, J. D., Opie, G. M., Ridding, M. C., & Semmler, J. G. (2013). A single bout of aerobic exercise promotes motor cortical neuroplasticity. Journal of Applied Physiology (Bethesda, Md. : 1985), 114(9), 1174–1182. https://doi.org/10.1152/japplphysiol.01378.2012

50. Michels, L., Martin, E., Klaver, P., Edden, R., Zelaya, F., Lythgoe, D., … O’Gorman, R. (2012). Frontal gaba levels change during working memory. PLoS ONE, 7(4). https://doi.org/10.1371/journal.pone.0031933

51. Mooney, R. A., Coxon, J. P., Cirillo, J., Glenny, H., Gant, N., & Byblow, W. D. (2016). Acute aerobic exercise modulates primary motor cortex inhibition. Experimental Brain Research. https://doi.org/10.1007/s00221-016-4767-5

52. Near, J., Ho, Y. C. L., Sandberg, K., Kumaragamage, C., & Blicher, J. U. (2014). Long-term reproducibility of GABA magnetic resonance spectroscopy. NeuroImage, 99, 191–196. https://doi.org/10.1016/j.neuroimage.2014.05.059

53. Okamoto, M., Dan, H., Sakamoto, K., Takeo, K., Shimizu, K., Kohno, S., … Dan, I. (2004). Three-dimensional probabilistic anatomical cranio-cerebral correlation via the international 10-20 system oriented for transcranial functional brain mapping. NeuroImage, 21(1), 99–111. https://doi.org/10.1016/j.neuroimage.2003.08.026

54. Pearce, A. J., Lum, J. A. G., Seth, S., Rafael, O., Hsu, C. M. K., Drury, H. G. K., & Tooley, G. A. (2014). Multiple bout rTMS on spatial working memory: A comparison study of two cortical areas. Biological Psychology, 100(1), 56–59. https://doi.org/10.1016/j.biopsycho.2014.05.002

55. Peirce, J. W. (2007). PsychoPy-Psychophysics software in Python. Journal of Neuroscience Methods. https://doi.org/10.1016/j.jneumeth.2006.11.017

56. Pontifex, M. B., Hillman, C. H., Fernhall, B. O., Thompson, K. M., & Valentini, T. A. (2009). The effect of acute aerobic and resistance exercise on working memory. Medicine & Science in Sports & Exercise, 41(4), 927–934. https://doi.org/10.1249/MSS.0b013e3181907d69

57. Ridding, M. C., & Ziemann, U. (2010). Determinants of the induction of cortical plasticity by non-invasive brain stimulation in healthy subjects. The Journal of Physiology, 588(Pt 13), 2291–2304. https://doi.org/10.1113/jphysiol.2010.190314

58. Rossi, S., Hallett, M., Rossini, P. M., & Pascual-Leone, A. (2009). Safety, ethical considerations, and application guidelines for the use of transcranial magnetic stimulation in clinical practice and research. Clinical Neurophysiology : Official Journal of the International Federation of Clinical Neurophysiology, 120(12), 2008–2039. https://doi.org/10.1016/j.clinph.2009.08.016

59. Rottschy, C., Langner, R., Dogan, I., Reetz, K., Laird, A. R., Schulz, J. B., … Eickhoff, S. B. (2012a). Modelling neural correlates of working memory: A coordinate-based meta-analysis. NeuroImage, 60(1), 830–846. https://doi.org/10.1016/j.neuroimage.2011.11.050

60. Rottschy, C., Langner, R., Dogan, I., Reetz, K., Laird, A. R., Schulz, J. B., … Eickhoff, S. B. (2012b). Modelling neural correlates of working memory: A coordinate-based meta-analysis. NeuroImage, 60, 830–846. https://doi.org/10.1016/j.neuroimage.2011.11.050

61. Rouder, J. N., Speckman, P. L., Sun, D., Morey, R. D., & Iverson, G. (2009). Bayesian t tests for accepting and rejecting the null hypothesis. Psychonomic Bulletin and Review, 16(2), 225–237. https://doi.org/10.3758/PBR.16.2.225

62. Silver, H., Feldman, P., Bilker, W., & Gur, R. C. (2003). Working memory deficit as a core neuropsychological dysfunction in schizophrenia. American Journal of Psychiatry, 160(10), 1809–1816. https://doi.org/10.1176/appi.ajp.160.10.1809

63. Smith, A. E., Goldsworthy, M. R., Wood, F. M., Olds, T. S., Garside, T., & Ridding, M. C. (2018). High-intensity Aerobic Exercise Blocks the Facilitation of iTBS-induced Plasticity in the Human Motor Cortex. Neuroscience, 373, 1–6. https://doi.org/10.1016/j.neuroscience.2017.12.034

64. Stagg, C. J., Bestmann, S., Constantinescu, A. O., Moreno Moreno, L., Allman, C., Mekle, R., … Rothwell, J. C. (2011). Relationship between physiological measures of excitability and levels of glutamate and GABA in the human motor cortex. Journal of Physiology, 589(23), 5845–5855. https://doi.org/10.1113/jphysiol.2011.216978

65. Stagg, C. J., Wylezinska, M., Matthews, P. M., Johansen-Berg, H., Jezzard, P., Rothwell, J. C., & Bestmann, S. (2009a). Neurochemical effects of theta burst stimulation as assessed by magnetic resonance spectroscopy. Journal of Neurophysiology, 101(6), 2872–2877. https://doi.org/10.1152/jn.91060.2008

66. Stagg, C. J., Wylezinska, M., Matthews, P. M., Johansen-Berg, H., Jezzard, P., Rothwell, J. C., & Bestmann, S. (2009b). Neurochemical effects of theta burst stimulation as assessed by magnetic resonance spectroscopy. Journal of Neurophysiology, 101(6), 2872–2877. https://doi.org/10.1152/jn.91060.2008

67. Stagg, Charlotte J. (2014, February 1). Magnetic Resonance Spectroscopy as a tool to study the role of GABA in motor-cortical plasticity. NeuroImage. Academic Press Inc. https://doi.org/10.1016/j.neuroimage.2013.01.009

68. Stagg, Charlotte J, Bachtiar, V., & Johansen-berg, H. (2011). What are we measuring with GABA magnetic resonance spectroscopy ? Communicative & Integrative Biology, (October), 573–575. https://doi.org/10.4161/cib.4.5.16213

69. Stavrinos, E. L., & Coxon, J. P. (2016). High-intensity Interval Exercise Promotes Motor Cortex Disinhibition and Early Motor Skill Consolidation. Journal of Cognitive Neuroscience, 1–12. https://doi.org/10.1162/jocn

70. Tanaka, H., Monahan, K. D., & Seals, D. R. (2001). Age-predicted maximal heart rate revisited. Journal of the American College of Cardiology. https://doi.org/10.1016/S0735-1097(00)01054-8

71. Thielscher, A., Opitz, A., & Windhoff, M. (2011). Impact of the gyral geometry on the electric field induced by transcranial magnetic stimulation. NeuroImage, 54(1), 234–243. https://doi.org/10.1016/j.neuroimage.2010.07.061

72. van Dongen, E. V., Kersten, I. H. P., Wagner, I. C., Morris, R. G. M., & Fernández, G. (2016). Physical Exercise Performed Four Hours after Learning Improves Memory Retention and Increases Hippocampal Pattern Similarity during Retrieval. Current Biology, 26(13), 1722–1727. https://doi.org/10.1016/j.cub.2016.04.071

73. Veen, J. W. van der, & Shen, J. (2013). Regional difference in GABA levels between medial prefrontal and occipital cortices. Journal of Magnetic Resonance Imaging, 38(3), 745–750. https://doi.org/10.1002/jmri.24009

74. Vidal-Piñeiro, D., Martín-Trias, P., Falcón, C., Bargalló, N., Clemente, I. C., Valls-Solé, J., … Bartrés-Faz, D. (2015). Neurochemical modulation in posteromedial default-mode network cortex induced by transcranial magnetic stimulation. Brain Stimulation, 8(5), 937–944. https://doi.org/10.1016/j.brs.2015.04.005

75. Voss, M. W., Vivar, C., Kramer, A. F., & van Praag, H. (2013). Bridging animal and human models of exercise-induced brain plasticity. Trends in Cognitive Sciences, 17(10), 525–544. https://doi.org/10.1016/j.tics.2013.08.001

76. Wang, J X, Rogers, L. M., Gross, E. Z., Ryals, A. J., Dokucu, M. E., Brandstatt, K. L., … Voss, J. L. (2014). Targeted enhancement of cortical hippocampal brain networks and associative memory. Science, 345(6200), 1054–1057. https://doi.org/10.1126/science.1252900

77. Wang, Jane X, Rogers, L. M., Gross, E. Z., Ryals, A. J., Dokucu, M. E., Brandstatt, K. L., … Voss, J. L. (2014a). Targeted enhancement of cortical-hippocampal brain networks and associative memory. Science, 345(6200), 1054–1057. https://doi.org/10.1126/science.1252900

78. Wang, Jane X, Rogers, L. M., Gross, E. Z., Ryals, A. J., Dokucu, M. E., Brandstatt, K. L., … Voss, J. L. (2014b). Targeted enhancement of cortical-hippocampal brain networks and associative memory. Science, 345(6200), 1054–1057.

79. Wang, Jane X, & Voss, J. L. (2015). Long-lasting enhancements of memory and hippocampal-cortical functional connectivity following multiple-day targeted noninvasive stimulation. Hippocampus, 25(8), 877–883.

80. Yamanaka, K., Yamagata, B., Tomioka, H., Kawasaki, S., & Mimura, M. (2010). Transcranial Magnetic Stimulation of the Parietal Cortex Facilitates Spatial Working Memory: Near-Infrared Spectroscopy Study. Cerebral Cortex, 20, 1037–1045. https://doi.org/10.1093/cercor/bhp163

81. Yoon, J. H., Grandelis, A., & Maddock, R. J. (2016). Dorsolateral Prefrontal Cortex GABA Concentration in Humans Predicts Working Memory Load Processing Capacity. The Journal of Neuroscience, 36(46), 11788–11794. https://doi.org/10.1523/JNEUROSCI.1970-16.2016

82. Ziemann, U., Paulus, W., Nitsche, M. A., Pascual-Leone, A., Byblow, W. D., Berardelli, A., … Rothwell, J. C. (2008). Consensus: Motor cortex plasticity protocols. Brain Stimulation, 1(3), 164–182. https://doi.org/10.1016/j.brs.2008.06.006

